# The MicrobeAtlas database: Global trends and insights into Earth’s microbial ecosystems

**DOI:** 10.1101/2025.07.18.665519

**Authors:** João Frederico Matias Rodrigues, Janko Tackmann, Lukas Malfertheiner, David Patsch, Eugenio Perez-Molphe-Montoya, Nicolas Näpflin, Daniela Gaio, Gregor Rot, Mihai Danaila, Matteo Eustachio Peluso, Marija Dmitrijeva, Thomas Sebastian Benedikt Schmidt, Christian von Mering

## Abstract

Environmental DNA sequencing has revolutionized our understanding of microbial diversity and ecology. Microbiomes have now been sequenced across the entire planet—from the deep subsurface to the mountain tops—covering a myriad of hosts, biomes, and conditions. Yet, the diversity of sequencing and processing strategies hampers universal insights.

MicrobeAtlas unifies more than two million microbiome samples in a single resource, harmonized to facilitate discoveries across technologies. Communities are hierarchically quantified at adjustable SSU rRNA marker gene resolution and feature detailed metadata, including rich geographic information. Connections to genome, phenotype, and ecological resources enable multimodal insights.

Microbial lineages can be reliably tracked across environments, including a ‘long tail’ of rare, uncharacterized species. Recurring community structures and geographic preferences become apparent, and global, taxonomy-specific generalism trends emerge. With MicrobeAtlas (www.microbeatlas.org), both known and newly described species and communities can readily be placed into ecological context, taking full advantage of earlier work.

## Introduction

The rapidly-evolving techniques of massively parallel sequencing have revolutionized microbiology. They have highlighted the enormous undescribed diversity of microbial species and provided valuable insights into microbial ecosystems^1–4^. The scale and breadth of sampled environments is increasingly advancing our understanding of microbial dispersal, niches and biogeography^5–7^. However, characterizing individual roles of microbial species within their communities and across geographic boundaries remains challenging.

The gold standard for characterizing microbial species requires isolation as pure cultures, followed by metabolic and physiological characterization. This often impractical and time-consuming approach is increasingly complemented by culture-independent methods that assemble microbial genomes directly from environmental sequences^2,8^. While these metagenome-assembled genomes (MAGs) enable first functional insights into uncultured microbes, their predictions do not always translate directly into phenotypes or ecological roles^9^. Furthermore, despite recent efforts^10–13^, the number of microbial species that have been cultured, phenotyped or (meta)genome-sequenced at sufficient quality is still orders of magnitude lower than the estimated total species richness^14,15^.

Marker gene sequencing offers an orthogonal approach to study microbes directly in their native community context. It systematically maps microbial species across environments, providing culture-independent insights into their abundance patterns, environmental conditions, and co-occurring taxa^6,7,16,17^. A cornerstone of this approach is the 16S small subunit ribosomal RNA gene (16S SSU rRNA), the most universally used molecular marker for microbial lineage identification. Large reference collections now aggregate millions of these sequences^18^, facilitating cross-study integration and revealing extensive microbial diversity. Despite limitations in quantifying lineage abundances^19^, SSU rRNA sequencing remains a cost-effective tool for microbiome analysis^20^. While phylogenetic resolution is constrained by sequence length and quality, advances in long-read technologies, such as circular-consensus and nanopore sequencing, are increasingly enabling species- and potentially even strain-level resolution^21,22^.

Previous marker gene studies, while restricted in scope, have proven instrumental in the characterization of the ecological preferences of microbial taxa^3,16,23^, the rare biosphere^24^, the identification of host-associated microbial taxa characteristic of health and disease^25–28^, and the identification of statistical signals indicative of biotic interactions^29–31^. However, many essential phenomena, including planetary biochemical cycling, global pathogen dispersal, and climate change, operate across environmental boundaries and thus require a global perspective.

Tracking microbes across multiple environments and studies remains challenging, as most existing large-scale datasets are confined to specific environments or hosts, and methods to systematically link these collections are lacking. The largest cross-environmental study to date, the Earth Microbiome Project, analyzed over 27,000 microbial community samples^7^, yet it captures only a small fraction of publicly available datasets and remains largely unintegrated with other studies.

Millions of environmental and clinical microbiome samples are hosted by repositories like the NCBI Sequence Read Archive (SRA^32^) and the EBI European Nucleotide Archive (ENA^33^), yet much of this raw data is not readily available for cross-study analyses. To help mitigate this issue, several databases and web resources have recently emerged to facilitate access to public microbiome data, typically focusing on single sequencing technologies. Some platforms analyze mainly metagenomic samples, providing high-resolution community profiles (SPIRE^34^, Sandpiper^35^), sometimes combined with functional insights from MAG assembly and gene- or protein-level characterizations (SPIRE^34^, MGnify^36^). A few specialize in specific environments, like the oceans (OMD^37^). However, due to the high cost of generating and analyzing metagenomic data, processed microbial communities remain limited to between tens of thousands and several hundred thousand per platform. Other resources (Qiita^38^, MGnify, IMNGS^39^) process subsets of the more abundant 16S amplicon data (up to half a million samples), but commonly applied *de novo* clustering or denoising techniques^40–43^ can lead to incompatibilities between studies analyzing different hypervariable regions of the 16S gene, hindering cross-study analyses within platforms. In addition, most resources focus on named taxa or fixed operational taxonomic units (OTUs), limiting the exploration of undescribed diversity at different taxonomic scales.

Here, we present MicrobeAtlas, an integrated resource of 2,390,937 analyzed microbial communities from all around the globe. We compared these microbiome samples against a reference database with over 1.5 million full-length 16S and 18S SSU rRNA sequences, pre-clustered hierarchically into hundreds of thousands of OTUs at selectable granularity. Mapping to a common, full-length reference enables OTUs and communities to be tracked across sequencing strategies (metagenomic, amplicon) and protocols (e.g., amplified 16S region) throughout the entire resource. Extracted metadata keywords and two-level environmental ontology assignments are available for each sample, complemented with carefully extracted geographic coordinates and spatial connections to terrestrial ecosystem definitions by the International Union for Conservation of Nature (IUCN^44^). Communities are hierarchically grouped into community clusters, providing a bird’s-eye view on global community patterns and potential niche distributions. Analyzing a robust subset of more than one million MicrobeAtlas samples reveals intriguing patterns: communities are broadly structured by environment, but also according to diverse, informative keywords related to specific conditions and hosts. Soil and aquatic microbiomes emerge as global richness hotspots with many poorly characterized members, contrasting most animal microbiomes. Individual OTUs can be reliably traced across the globe, revealing detailed habitat generalism and specialism trends, as well as geographic preferences. This includes the uncharted majority of OTUs without cultured representatives, sequenced genomes, or robust taxonomic assignments, revealing first glimpses of their ecology. Finally, we identified blind spots in current sampling efforts that point to geographic regions, ecological lifestyles, and specialized biomes that require more concerted attention by the community.

## Results

### The MicrobeAtlas dataset

We comprehensively screened the NCBI SRA database for microbial community sequencing samples and downloaded raw reads and metadata for 2,390,937 samples (52,950 studies). Utilizing a sequencing strategy-agnostic workflow, we quality-checked and mapped 6.87 trillion raw reads against a custom database of 1,536,850 full-length 16S/18S marker gene sequences. After removing samples with low community complexity, coverage, and sequencing depth (Methods), we obtained a dataset of 1,153,349 consistent, comparable community abundance profiles (24,349 studies) used for analyses in this study (Fig. 1).

**Figure 1.**
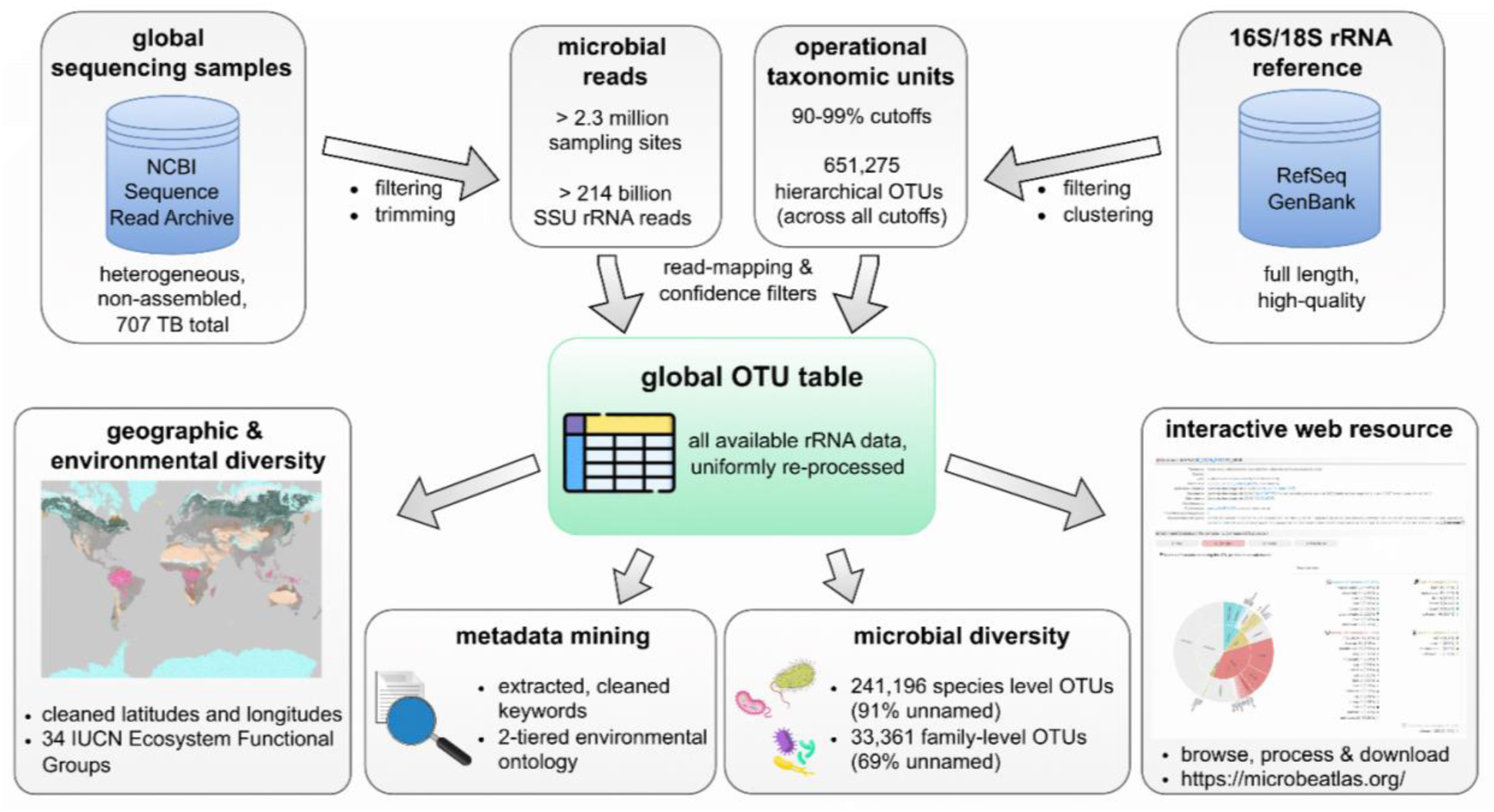
MicrobeAtlas data retrieval and analysis pipeline. Raw data from targeted and untargeted microbial sequencing were downloaded from the NCBI Sequence Read Archive, cleaned, and mapped against OTUs pre-clustered at flexible thresholds (e.g., species-level: 98%, family-level: 90%). The resulting OTU table was crosslinked with geographic information, consensus taxonomies, and community cluster information, and detailed ontological and keyword-based metadata. An interactive web application allows in-depth explorative analysis and download of whole communities and individual taxa.

How different community members are defined and distinguished can profoundly impact ecological analyses. In order to enable flexible definitions, we therefore clustered our reference sequences into hierarchical OTUs with 90% to 99% sequence identity (33,361 to 241,196 OTUs, default: 97%-OTUs) and profiled the communities of all samples at each OTU level. In order to reduce redundancy and facilitate niche-focused analyses, we additionally hierarchically clustered community profiles into 356,182 microbial community clusters, each potentially representing an ecological niche with distinctive microbial composition.

A key component of MicrobeAtlas is its rich metadata. For this purpose, we extracted information on environments, geo-locations, and informative keywords from raw metadata to provide detailed biological and technological context for each community. We categorized samples into a simple, two-tiered environmental ontology (Supplementary Table 1) and, where possible, additionally linked terrestrial samples geographically to the IUCN Global Ecosystem Typology, providing more precise ecosystem definitions based on macro-ecological characteristics (Supplementary Table 2 and 3). The resulting dataset spans four main environments (animal, aquatic, soil, plant) and 65 sub-environments, covering a plethora of habitats, hosts, and conditions from all continents. In addition, we linked MicrobeAtlas OTUs to culture collections (BacDive^45^) and genome clusters (proGenomes3^46^), connecting ecological distributions with phenotypic and genetic information. To our knowledge, MicrobeAtlas houses the largest and most diverse collection of cross-study comparable microbial community profiles available to date.

To enable intuitive exploration of this dataset, we created a feature-rich website (www.microbeatlas.org) that allows the tracking, comparison, and download of studies, samples, and individual taxa from around the globe (Fig. 1). OTUs can be searched based on taxonomic or metadata matches and can be visualized on a global map, together with their abundance distributions across environments. Similarly, samples and projects can be identified using rich metadata search and explored in terms of community composition, keywords, and technical information, across flexible OTU definitions.

### A map of global microbiome structure

We projected 26,487 robust community clusters (with at least 5 samples per cluster) to obtain a global view of planetary microbial diversity across environments (Fig. 2a). While a few large projects individually contribute large numbers of samples (Supplementary Fig. 1), smaller studies collectively accounted for the vast majority of community clusters (92.6%, Fig. 2b), a proxy for the breadth of the captured community compositions. We found that samples from selected major consortium studies typically had close matches within smaller studies, but in contrast samples from smaller studies were more distant from major studies (min. Bray-Curtis dissimilarity: 0.31 vs. 0.67; Supplementary Fig. 2), further emphasizing the additional variation contributed by smaller studies. Community clusters were mainly animal-associated (63.5%, human 23.0%), followed by aquatic (19.3%, marine 4.3%), soil (14.3%, agricultural 2.8%), and plant (2.9%, rhizosphere 0.6%) (Supplementary Fig. 3). The number of community clusters did not saturate with increasing sample size, suggesting a large fraction of thus-far undiscovered community clusters remains in each main environment (Fig. 2d). In contrast, OTU richness rarefaction curves were closer to saturation, indicating that the majority of reference OTUs has been detected within their respective environments (with the notable exception of plant communities, Fig. 2d). Notably, this saturation is constrained to our reference database and therefore conservative (Supplementary Fig. 4).

**Figure 2.**
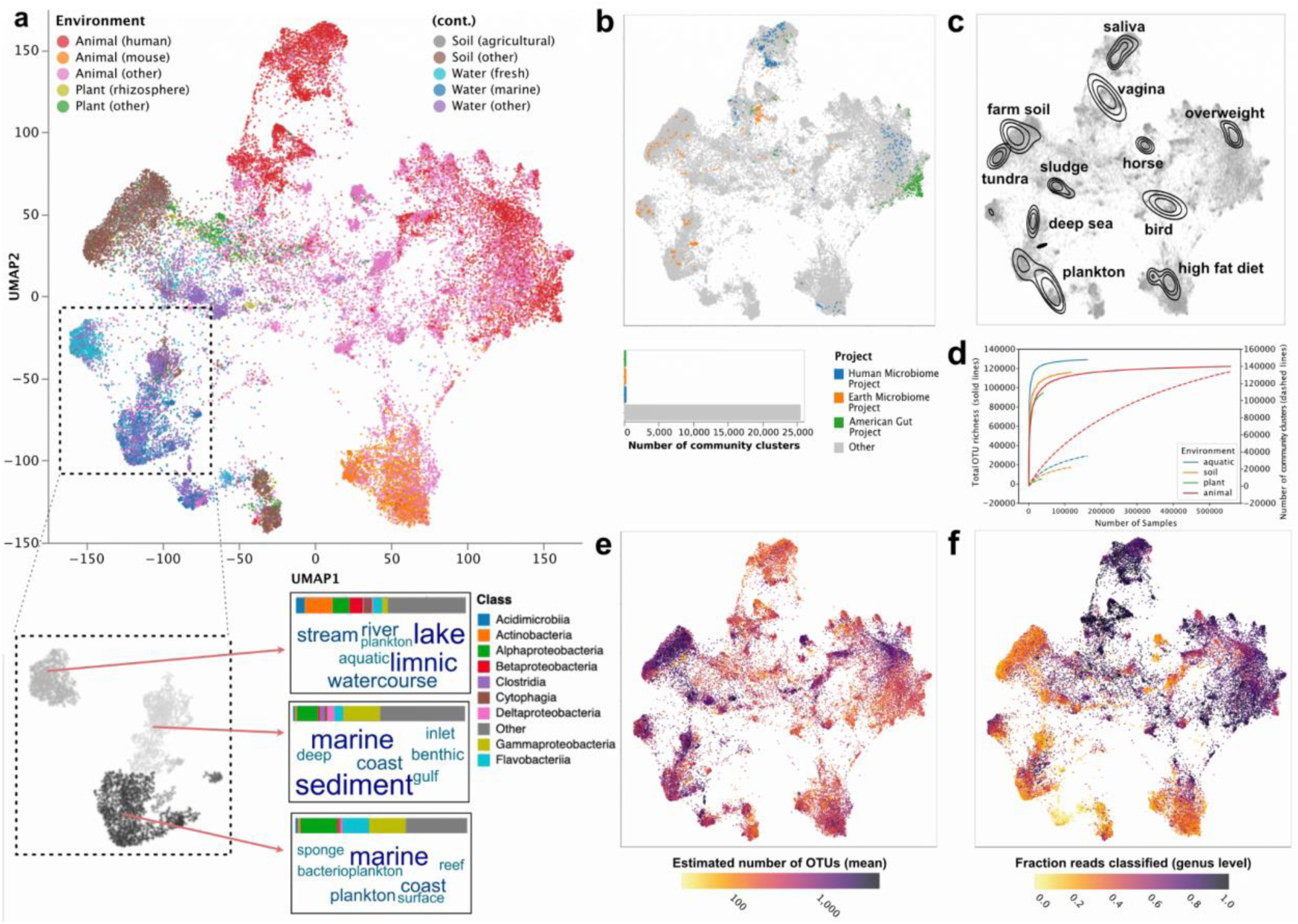
UMAP projections of 26,487 robust community clusters, derived from 1,153,349 filtered sequencing samples. **a**, Top: community clusters (dots) based on Bray-Curtis dissimilarities, colored according to consensus main environment and sub-environment (“other” includes clusters without sub-environment annotation). Clusters with no confident main environment were removed. Bottom: grayscale colors depict DBSCAN density clusters computed from UMAP coordinates (euclidean distance). Class-level bar charts represent average relative abundances per taxonomic class and DBSCAN cluster (normalized per cluster). Word clouds show the seven most prevalent keywords per DBSCAN cluster, text size and darkness according to keyword frequency. **b**, Top: community clusters colored by major sequencing projects (HMP, EMP, American gut) or smaller studies (other). Bottom: number of community clusters per project type. **c**, Kernel density estimates of example keyword frequencies across community clusters (normalized per keyword). **d**, The number of total community clusters (dashed lines) or OTUs (solid lines, OTU level: 97%) as a function of sampling effort (number of samples), stratified by sample environment. **e-f**, Community clusters colored by estimated OTU richness (e, Methods, OTU level: 97%) or fraction of reads classified at genus level (f).

The projection revealed distinct, biologically informative patterns: clusters grouped by main and sub-environment (Fig. 2a, top; PERMANOVA (unifrac-w): R^2^ = 14.7% (main), R^2^ = 10.6% (sub), p = 0.001 for both), with environmental (aquatic, soil) and plant clusters typically separated from animal-associated clusters. Most freshwater samples are located between marine and soil clusters (supported by diverse dissimilarity metrics, Supplementary Fig. 5a), indicating potential microbial exchange between soil and freshwater environments, in line with previous reports^47,48^. Similarly, environmental samples are closer to surface body sites (skin, oral, nasal/airways, vagina) than to gut communities (Supplementary Fig. 5b), likely reflecting higher prevalence of environmental microbes on more accessible body sites.

Among larger projection clusters, we observed detailed patterns, where neighboring yet distinct sub-clusters differ in both microbial composition and metadata annotation in biologically interpretable ways. Using keyword analysis, we characterized a freshwater projection cluster in more detail and distinguished it from two marine clusters, one related to benthic and sediment communities, and the other dominated by planktonic and host-associated communities (Fig. 2a, bottom). These clusters also differ in community composition at the class level, with marine clusters being more similar to each other than to the freshwater cluster (class level Bray-Curtis: 0.25 vs. 0.42). In addition, we found examples of highly localized keywords (Fig. 2c) distinguishing projection regions enriched for certain soil types (tundra, farm soil), hosts (horse, bird), or conditions (high-fat diet). A more targeted extraction of host information via host-specific fields and taxonomic IDs recovered 2682 host species, the most frequent ones often localized in specific regions of the projection (Supplementary Fig. 6). Mouse was the second most frequent host (after human, Supplementary Fig. 7) and, notably, mouse gut communities formed a separate cluster in our projection (Fig. 2a, Supplementary Fig. 8). Closer examination of semantic keyword groups (Methods) revealed enrichment of categories related to (experimental) perturbation and laboratory conditions in this cluster, including terms such as “antibiotics”, “disruption”, “heat-killed”, “dose-dependent”, “stress” (Supplementary Table 4). In contrast, the core human gut cluster was closer to non-mouse hosts in the projection and featured more observational keyword groups related to subject traits or diseases, such as “female”, “twin”, “infant”, “malnutrition”, “ulcer” (Supplementary Table 5, Supplementary Fig. 8).

In line with previous studies, we found that environments differ strongly in diversity. Estimated mean OTU richness per community cluster (Fig. 2e) was particularly high in environmental (soil, freshwater, marine; mean 715-1257 OTUs), waste water (1142 OTUs), and ruminant animal samples (988 OTUs), independent of sequencing technology (Supplementary Fig. 9). In contrast, we found lower richness in other animals (mean 490 OTUs; in particular insects with 281 OTUs) and surface plant organs (539 OTUs). Counting observed OTUs after rarefaction led to the same qualitative conclusions (Supplementary Fig. 10). In terms of taxonomic coverage, human samples had high fractions of reads classified at least to genus level (mean 78.4%; Fig. 2f), while environmental communities had strongly reduced fractions (down to mean 35%), indicating notable under-characterization of these high-richness habitats. This was particularly apparent for soils, sediments, and marine samples (37.2%, 35.9%), but also freshwater (45.7%), and, intriguingly, mouse communities (42.7%) (Supplementary Fig. 11). In general, classification fraction and richness were negatively correlated across community clusters (ρ = −0.36, p < 1 × 10^-100^; Supplementary Fig. 12).

Finally, we also identified technology-associated effects. A minor region of the projection, comprising community clusters from diverse environments, was strongly enriched for Eukaryotes (mean Eukaryotic fraction: 88.5%, Supplementary Fig. 13), particularly Fungi (29.3%) and Metazoa (17.5%). Samples in a manually inspected subset were typically generated using Eukaryote-specific 18S primers or, in some cases, by amplifying 16S genes from Eukaryotic organelles, particularly mitochondria and chloroplasts. Beyond Eukaryote-specific patterns, we also observed signs of localized clustering among metagenomic samples (i.e. generated using whole-genome shotgun protocols, Supplementary Fig. 14a) compared to amplicon samples (Supplementary Fig. 14b). However, these effects were small compared to the main environment signals described above (PERMANOVA (unifrac-w): R^2^ = 0.6%, p = 0.001).

### Ecological insights into the uncharacterized microbial majority

BacDive and proGenomes3 are two of the most comprehensive collections for culture strain and genome information, respectively. We mapped these databases to MicrobeAtlas OTUs and found that most available strains (92.6%) and genomes (97.3%) were covered. In contrast, the vast majority of OTUs in MicrobeAtlas (n=99,241, 88.7%) lacked cultured isolates, genomes, and detailed taxonomy (Fig. 3a-c), making them potential representatives of “microbial dark matter”^49^. As expected, OTUs without any culture strain or genome information (ncg-OTUs) tended to be less taxonomically classified: only 21.6% had confident consensus assignments at the genus level (class: 83.0%), whereas OTUs with culture strain or genome annotations (cg-OTUs) reached 65.4% (class: 93.0%) (Fig. 3b). Notably, 14,985 ncg-OTUs were classified only at the phylum level or higher, outnumbering all cg-OTUs combined (12,543). Among them, 4,988 ncg-OTUs (5%) lacked any consensus phylum annotation and may potentially represent uncharacterized phyla. In addition, the vast majority of described (candidate) phyla were dominated by ncg-OTUs, while only around half of the described genera showed this trend (Fig. 3c).

**Figure 3.**
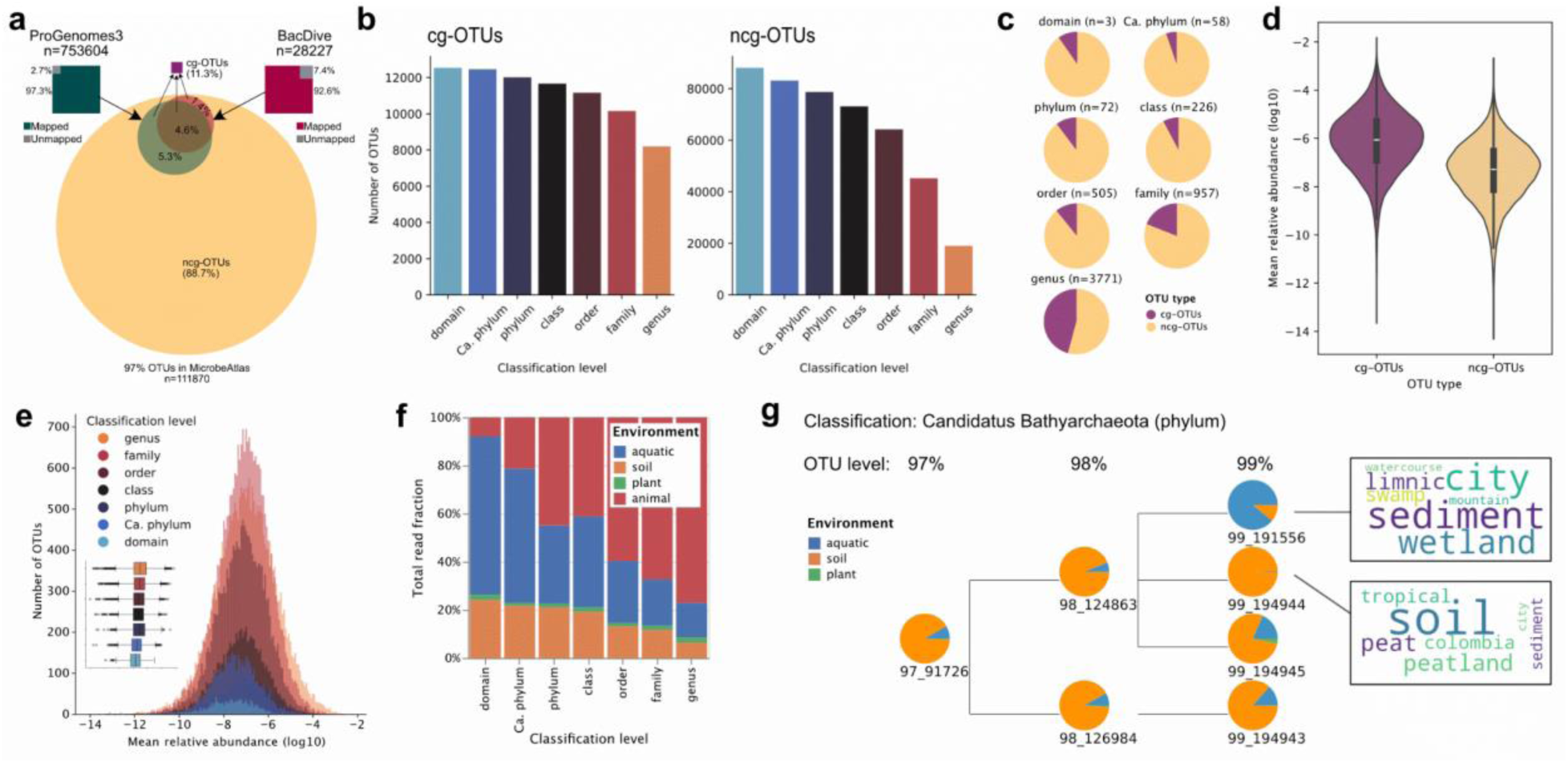
Taxonomic characterization and ecology of MicrobeAtlas OTUs. **a**, Circles show MicrobeAtlas OTUs that were mapped (green and red circle) or not mapped (yellow circle) to proGenomes3 or BacDive sequences, counting best hits with >97% identity. Mapped OTUs are collectively referred to as culture/genome-OTUs (cg-OTUs), while unmapped OTUs are referred to as non-cg-OTUs (ncg-OTUs). Squares show the fraction of proGenomes3 (green) and BacDive (red) sequences mapped to MicrobeAtlas OTUs. Unmapped sequences are shown in grey. **b**, Numbers of OTUs classifiable at different taxonomic ranks (blue: less classified, red: more classified), with and without culture strain or genome annotations. **c**, Fractions of taxa with at least one annotated culture strain or genome (purple) at each taxonomic rank. **d**, Mean relative abundances (log10) of species with and without culture strain or genome annotations. **e**, Abundance distributions of OTUs classifiable at different taxonomic ranks (blue: less characterized). Inset: the same abundances depicted as box-plots (box: interquartile range, whiskers: 95% confidence interval). **f**, Environmental distributions of OTUs classifiable at different taxonomic ranks (fraction of mapped reads, normalized per rank; Eukaryotes removed). **g**, Environmental distributions of an uncharacterized 97%-OTU and its child OTUs (mean relative abundances, re-scaled). Word clouds show the seven most prevalent keywords for two OTUs across inhabited samples, weighted by relative abundance. Text size according to abundance-weighted keyword frequency.

MicrobeAtlas allowed us to globally track both taxonomically characterized and uncharacterized OTUs to gain insights into their ecology. We observed a notable number of ncg-OTUs among the most abundant microbes across environments (17 out of the top 150), despite cg-OTUs being on average 16 times more abundant than ncg-OTUs (Fig. 3d). Top ncg-OTUs included many Bacteroidales OTUs (e.g., Muribaculaceae, *Bacteroides*; Supplementary Table 6), but also an archaeal OTU classified as *Candidatus* Nitrosocosmicus, detected in 82,660 samples (4,232 studies) across all continents (Supplementary Fig. 15) and most prevalent in soil (56%). Notably, several OTUs without phylum annotations also reached high abundances (Fig. 3e, up to the 97th percentile), for instance OTU 97_17665 with 16,222 samples (2,736 studies), most abundant in drinking water-associated communities and biofilms (Supplementary Fig. 16).

Out of 99 bacterial and archaeal phyla, 43 featured exclusively phylum-level OTUs, and only a minority of phyla were on average classified at the family or genus level (26 and 2 phyla, respectively; Supplementary Fig. 17c). In general, we found classification level, phylum diversity (number of distinct 97% OTUs per phylum) and global abundance to be significantly correlated: well-classified phyla tended to be OTU-rich and abundant (ρ = 0.33, *P* < 0.001, and ρ = 0.67, *P* < 1 × 10⁻¹⁰, respectively). Nonetheless, we also observed numerous phyla classified only at the phylum level that were highly diverse (hundreds of 97% OTUs) and abundant (e.g., *Candidatus* Marinimicrobia, *Candidatus* Absconditabacteria; Supplementary Fig. 17b).

When investigating microbial habitats, we found that less characterized OTUs tended to be more common in environmental samples (soil, aquatic, plant) (Fig. 3f): while only 20.8% of reads of OTUs classified at genus-level mapped to environmental communities, this number increased to 57.0% at class-level and, strikingly, to >77% at candidate phylum level and above. This trend was strongest for domain-level OTUs, the least characterized group, in aquatic environments: 65.8% of domain-level reads mapped to aquatic communities (up from 14.3% for genus-level OTUs), most frequently to the sub-environments marine, sediment, and waste water (Supplementary Fig. 18).

Each OTU in MicrobeAtlas is part of a nested hierarchy generated by agglomerative clustering of 16S sequences (Fig. 1). This structure allows exploration of microbial lineages at varying sequence identity thresholds (e.g., 97–99%) and enables coarse-grained eco-evolutionary analyses of largely uncharacterized OTUs (Fig. 3g). For instance, we found an example of notable environmental separation within a 97%-OTU classified only as *Candidatus* Bathyarchaeota (Fig. 3g). This OTU and most of its children were predominantly present in soil-like environments, with minor occurrences in aquatic samples. However, one child OTU at the 99% level (99_191556, found in 314 samples from 171 studies) specialized towards predominantly aquatic environments, while one of its sister OTUs (99_194944, 474 samples, 165 studies) was encountered almost exclusively in soils. Comparative keyword analysis indicated 99_191556 to be more abundant in limnic and wetland environments, while 99_194944 seems to thrive in peaty soils (Fig. 3g).

### Biogeographic trends across continents and biomes

The extent of MicrobeAtlas enables detailed biogeographic analyses of single taxa and whole microbiomes around the globe. All continents and climate zones are represented, from the equator to the poles (Fig. 4a). Still, we observed pronounced geographic sampling biases, where the Northern Mid Latitude has been studied extensively (hot spots in Europe, North America, and East Asia, Fig. 4a, Supplementary Fig. 19a), with a major focus on animal microbiomes (Supplementary Fig. 19b). In contrast, lowly populated areas (polar regions, sub-arctic boreal regions, deserts) and the Southern hemisphere are comparatively undersampled (Fig. 4a, Supplementary Fig. 20).

**Figure 4.**
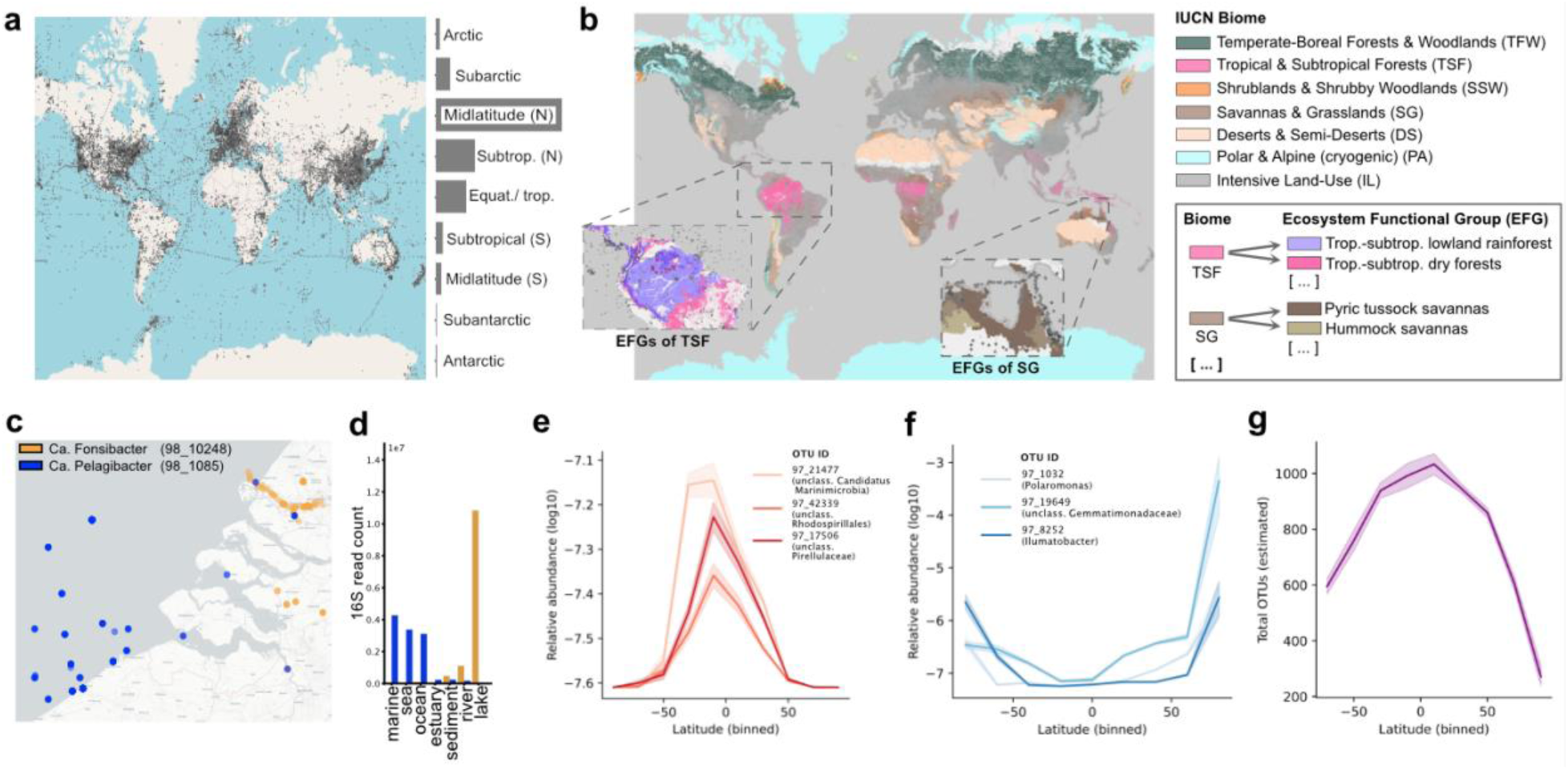
Biogeographic patterns of microbial taxa and communities. **a**, Global distribution of microbial sequencing samples across climate zones. **b**, IUCN Biome and Realm maps, relating microbial communities to macro-ecological features. Gray dots within magnified areas depict MicrobeAtlas samples. **c**, Example of niche separation between two Pelagibacterales OTUs (dots are observed occurrences). **d**, Global read counts by sub-environment for each OTU in (c). **e-f**, Example OTUs with high preference for tropical and subtropical regions (e) or polar and subpolar regions (f). Lines and error bands represent means and 95% confidence intervals of log-transformed relative abundances (1000 bootstraps) within 10 equally-sized latitudinal bins. **g**, Estimated OTU richness (Methods) of marine communities across latitudes. Lines, error bands, and bins defined as in panels e-f.

The integration of IUCN hierarchical ecosystem definitions enabled us to comprehensively compare microbiomes between distinct macro-habitats across scales, defined in terms of function, vegetation type or anthropogenic pressures (Fig. 4b). MicrobeAtlas features diverse microbiomes from all seven terrestrial IUCN Biomes, as well as 34 Ecosystem Functional Groups (EFGs), ranging from tropical forests and grasslands to deserts and cryogenic zones. We found that the majority of the 65,909 IUCN-mappable terrestrial samples originate from heavily anthropogenic habitats (“Intensive Land-Use” biome (75.7%), in particular the EFG “Urban and industrial ecosystems” (17.6%)), followed by the biomes “Savannas and Grasslands” (9.6%) and “Temperate-boreal Forests and Woodlands” (7.7%) (Supplementary Fig. 20, Supplementary Table 2 and 3). In contrast, the ten least-sampled EFGs (less than 10 samples each) are spread across multiple Biomes and include “Ice sheets, glaciers and perpetual snowfields”, and multiple EFGs related to heathlands, tropical or alpine environments, and every type of savanna (Supplementary Fig. 20b). Notably, six of these had no mapped samples (e.g., “Cool temperate heathlands”, “Hummock Savannas”). A less stringent keyword-based search recovered more samples from underrepresented environments (e.g., 5638 samples for “glacial” and 20 samples for “hummock”); these however lacked robust geographic links to IUCN-defined ecoregions.

When investigating biogeographic trends at the OTU level, we found striking examples of global niche separation. For instance, two OTUs classified as *Candidatus* Pelagibacter and *Candidatus* Fonsibacter, respectively (both Pelagibacterales), are globally prevalent in aquatic environments and occasionally geographically co-localized (Fig. 4c). Yet, they show clear environmental distinction where Pelagibacter is widely distributed across global oceans while Fonsibacter is almost exclusively found in freshwater environments, in particular lakes (Fig. 4d).

In terms of broader biogeographic trends, we identified 37,393 97%-OTUs with significant latitudinal preferences in at least one environment: 93.3% of these were more abundant around the equator (e.g., unclassified Rhodospirillales and *Candidatus* Marinimicrobia OTUs, Fig. 4e) and the remainder more abundant in polar regions (e.g., *Polaromonas* and *Ilumatobacter* OTUs; Fig. 4f). We observed most latitudinal trends in aquatic samples (54% of significant correlations), followed by animal (31%) and soil (14%), and very little in plants (2%, possibly due to undersampling). Notably, soil samples harbored both equatorial- and polar-associated microbes (63% equatorial), in contrast to all other environments (86% − 99% equatorial correlations). In line with high equatorial associations in aquatic OTUs (97%), marine OTU richness also peaked near the equator (max. richness bin: 0°N to 20°N, Fig. 4g), consistent with latitudinal species diversity gradients observed for many multicellular organisms^50^ and matching previous reports on marine microbes^3^. Interestingly, we observed comparable diversity spikes in sufficiently sampled terrestrial biomes (max. richness bin: 20°N to 40°N, biomes: TFW, DS, IL; Fig. 4b, Supplementary Fig. 21), despite soil OTUs showing fewer equatorial associations individually than marine OTUs. Counting observed OTUs after rarefaction led to the same qualitative conclusions, both for marine and terrestrial habitats (Supplementary Fig. 22, Supplementary Fig. 23).

### Environmental and taxonomic generalism patterns

Habitat generalism is an important ecological axis: while some species specialize in particular environmental niches (habitat specialists), others thrive in diverse environments and conditions (habitat generalists). To investigate global habitat generalism trends, we devised a score that globally quantifies habitat generalism tendency based on per-environment abundances at a coarse-grained level. Under this entropy-based metric (Methods), OTUs with even abundance distributions across major environments (aquatic, soil, plant, animal) score closer to one (i.e. are more generalist), while species with uneven distributions, i.e. with abundance peaks at one or few environments, score closer to zero (are more specialist) (Fig. 5a).

**Figure 5:**
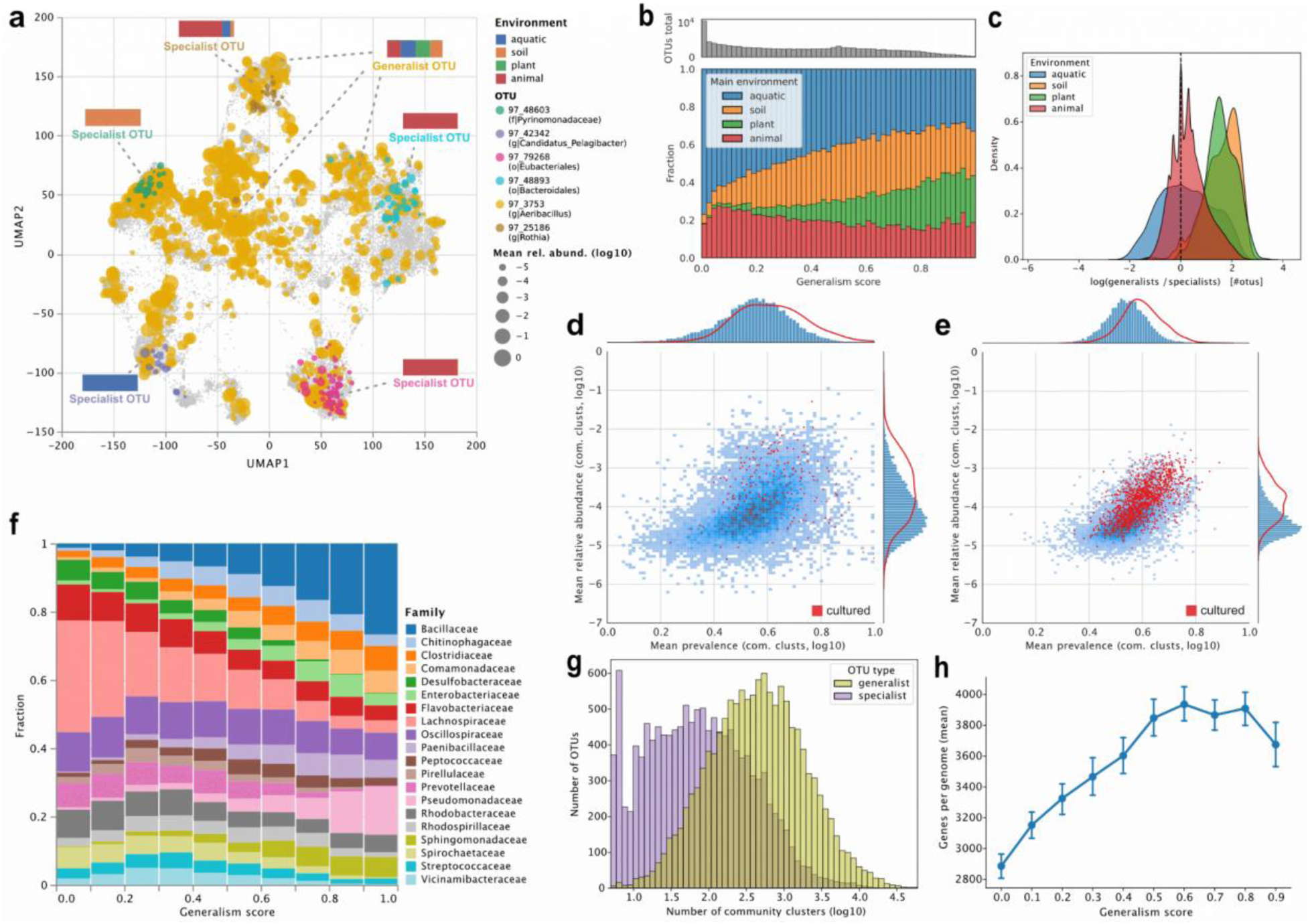
Habitat generalism and specialism trends across environments and taxonomy. **a**, UMAP community cluster abundance distributions of one generalist (yellow circles) and five specialist OTUs (other colors; gray: no displayed OTU present). Bar charts: environmental relative abundance distributions per OTU. **b**, Top histogram: number of OTUs in each normalized generalism score bin. Bottom bar charts: fraction of OTUs in each generalism score bin, stratified by preferred environment (defined by highest relative abundance). **c**, Log-ratios of the number of top-10% generalists to the number of top-10% specialists for each sample, grouped by environment. **d-e**, Relationship between average prevalence and relative abundance (zeros removed) across community clusters, separated by top-10% specialists (d) and top-10% generalists (e). Red dots: OTUs with cultured representatives. Marginal densities for cultured representatives (red lines) are kernel density estimates. **f**, Fractions of the 20 most frequent taxonomic families, binned by normalized generalism score. OTUs not mapping to any family were excluded. **g**, Density of the number of community clusters (log10) across top-10% generalist (green) and top-10% specialist (purple) OTUs. **h**, Average number of genes across proGenomes3-mappable OTUs within each generalism score bin. Lines and error bands represent means and 95% confidence intervals (1000 bootstraps).

On this level of environmental granularity, we found that OTUs show a clear tendency towards habitat specialism (39.6% of OTUs have scores in the bottom quarter of the range, but only 10.3% lie in the top quarter; Fig. 5b, top histogram), mostly thriving in aquatic and, to a lesser extent, animal habitats (Fig. 5b, bottom stacked bar charts). In contrast, OTUs abundant in soil and plant-associated environments appeared more generalist, i.e. were more likely to also be abundant in animal and aquatic environments. This pattern remained when adjusting for potential assignment ambiguities between plant rhizosphere and soil (recomputing generalism scores without the “plant” category, Supplementary Fig. 24). Similarly, soil and plant communities on average harbored significantly more generalist species than aquatic or animal communities (mean log ratio 1.58 vs. 0.30, p < 10^-99^, Mann-Whitney U test; Fig. 5c), again robust to potential soil/rhizosphere ambiguities (Supplementary Fig. 25). Certain generalists reached scores close to the theoretical maximum (e.g., OTU 97_3808 (genus: *Gordonia*) with 0.9991), indicating highly even abundance distributions across environments.

On average, top generalists inhabit significantly more community clusters than top specialists (1239.5 vs 263.9, p < 10^-99^, Mann-Whitney U test; Fig. 5g), indicating wider dispersal overall. However, within community clusters (proxies for occupied niches), highly generalist and specialist OTUs (top and bottom 10%) both had similar mean abundances (4.8 × 10^-5^ vs. 9.2 × 10^-5^) and prevalences (0.546 vs. 0.549) (Fig. 5d, e), indicating comparable colonization success within these realized niches. Notably, specialist OTUs exhibited greater variation in both metrics: some specialists dominated their niches more than comparable generalists, while other specialists showed the opposite pattern (Fig. 5d, e). In addition, we found clear signals of culturing biases: generalists were significantly more likely to have cultured representatives than specialists (odds ratio = 6.85, p < 10^-99^, Chi-squared test; Fig. 5d, e) and, in addition, more abundant and prevalent species were also significantly more likely to have been cultured (AUC = 0.72 and 0.66, p < 10^-99^ both, Mann-Whitney U test).

In terms of taxonomy, we observed notable family-level shifts across the generalism gradient (Fig. 5f). Specialist OTUs cover known host-associated families, most notably Lachnospiraceae and Prevotellaceae, but also environmental (in particular aquatic) groups such as Desulfobacteraceae. In contrast, high-scoring generalists include families known for their adaptability and dispersal potential, most notably Pseudomonadaceae and Bacillaceae. Despite these trends, each investigated family harbored OTUs from both sides of the generalism spectrum and some families showed a remarkably balanced distribution of generalists and specialists (e.g., Oscillospiraceae, Flavobacteriaceae).

Generalists may differ from specialists in terms of genomic characteristics (e.g., genome size, number of genes), potentially providing increased flexibility in the face of varying environmental pressures. Indeed, when we investigated the average number of genes per genome within each OTU (available for 7427 97% OTUs, based on our 16S-based mapping to proGenomes3), we found that gene counts increased consistently with higher generalism scores. Counts peaked for OTUs with generalism scores between 0.6 and 0.7, averaging 3935.6 genes (mean), a significant 36.4% increase compared to OTUs with generalism scores between 0.0 and 0.1 (p = 10^-46^, Mann-Whitney U test). In line with earlier observations (Fig. 5b, c), plant- and soil-associated OTUs in general tended to have larger genomes than animal and aquatic OTUs (Supplementary Fig. 26). To account for this, we sub-sampled each generalism bin for equal environmental representation, resulting in a similar gene count increase that, however, plateaued earlier and reached lower statistical significance (32.7% increase, p = 10⁻⁴; Supplementary Fig. 27).

## Discussion

Here, we present an initial analysis of MicrobeAtlas, featuring 2,390,937 processed samples—the largest collection of cross-study compatible microbial community profiles to date. The vast majority of community-level diversity in MicrobeAtlas comes from smaller studies, underscoring the value of integrating data beyond major consortium efforts. We complemented the dataset with systematically extracted and processed metadata information, combined with links to genome and strain collections. The entire database, including community profiles, OTU-level data, and metadata, is available as a comprehensive package, downloadable and interactively explorable on our feature-rich website at www.microbeatlas.org.

In our global analysis, we found both expected and new ecological patterns of interest. The notable fine-structure we observed among major environmental community clusters indicates rich, partially undiscovered sources of microbial variation. While we identified potential contributors—such as detailed sub-environmental patterns and diverse host-related structures— these examples merely scratch the surface. Nonetheless, they illustrate the depth and explorative potential of MicrobeAtlas. Importantly, these patterns were initially identified through keyword frequency analysis and semantic clustering, which we see as powerful explorative tools to detect potential community drivers within non-standardized metadata. As one example, a distinct mouse gut cluster showed low taxonomic resolution, consistent with prior reports of under-characterized microbiota in both lab and wild mice^51^. Frequent experiment-related terms and semantic clusters suggest lab conditions may contribute to this separation. These microbiomes also differ markedly from human gut communities, highlighting ongoing concerns about the relevance of mouse models^52,53^.

Biogeography is a major focus of MicrobeAtlas, supported by automatically extracted, cleaned coordinates and IUCN ecosystem annotations. The wide-spread latitudinal trends we detected across taxa include evidence for (sub-)tropical diversity hotspots in marine and terrestrial environments, expanding on earlier reports^3,6^ and contributing to long-standing discussions of latitudinal diversity gradients in macro-ecology^50,54^. Indications of sharp niche separation between two geographically co-localized species^55^, abundant in freshwater and marine coastal areas, respectively, further illustrate the breadth of geographic analysis avenues.

Our quantification of high-level habitat generalism covers hundreds of thousands of OTUs across more than one million communities, providing a useful resource for the research community. The strong generalism-specialism segregation we observed among taxonomic families coincides with established host associations (e.g., Lachnospiraceae^56^) and resilient, broadly adaptable groups (e.g., Pseudomonadaceae^57^, Bacillaceae^58^) at opposite sides of the spectrum. Notably, Enterobacteriaceae—harboring notorious pathogens such as Enterobacter, Salmonella, and Klebsiella—exhibit concerningly high scores, suggestive of broad environmental reservoirs^59,60^. The enrichment of generalists in soils and plants, along with their tendency towards larger genomes, further indicates enhanced environmental robustness and dispersal potential of these communities, possibly facilitated by expanded genetic repertoires^61^.

Our investigation of global microbiomes revealed notable dark spots in global study efforts. The vast majority of MicrobeAtlas OTUs remains largely uncharacterized, lacking cultures, genomes, and detailed taxonomy. Among them, we identified numerous highly abundant yet understudied OTUs thriving in environmental samples, emphasizing the need for further microbial characterization in global soils, lakes, rivers, and oceans. We found culturing biases to particularly favor habitat generalists and abundant OTUs, stressing the importance of high-throughput culturing efforts targeting a broader microbial spectrum^62^—critical, for instance, in bioactive compound discovery^37^. With keyword-based metadata, aided by web-based exploration and easy data extraction for co-occurrence and other analyses, MicrobeAtlas offers data-driven ways to prioritize these enigmatic yet likely impactful taxa.

Major gaps also persist at the environmental and geographic levels. Animal microbiomes are notably over-represented, making up over 50% of all samples, whereas marine samples, for instance, account for less than 8%, despite oceans covering the majority of Earth. While animal studies are of clinical importance, understanding global environment-critical phenomena such as biogeochemical cycles and climate change requires similar characterization efforts for environmental communities^63^. We found similar sampling biases across (bio)geographic axes: while sampling epicenters in North America, Europe, and East Asia become apparent, other regions are notably under-represented, including most climate zones (in particular tropical and polar zones), ecosystems (e.g., alpine regions, savannas), and the Southern hemisphere in general. Exploring understudied environments, regions, and traditional cultural practices continues to reveal new biological insights^49,64^, underscoring the importance of targeted sampling efforts in these blind spots.

In the future, MicrobeAtlas will continue to receive regular updates, with the upcoming release scheduled to comprise 6.3 million samples, currently in the final stages of processing. Looking forward, the resource will benefit from the extension of SSU rRNA reference collections, aided by long-read sequencing technologies^22,65^. Emerging machine learning tools that extract information from noisy metadata^66^ should further improve sample categorization, complementing increasingly consistent metadata standards^67^. Cross-linking MicrobeAtlas to Earth observation data would provide an intriguing interdisciplinary avenue to increase data richness and extend global prediction models with microbial information^68^. We are only beginning to unravel the intricate network of global microbial ecosystems, but data-driven resources such as MicrobeAtlas will be key to advancing that goal.

## Methods

### Sample selection and download

Metadata summary files for all NCBI Sequence Read Archive (SRA) samples were downloaded from the NCBI ftp server^69^. Every sequencing run from projects with metadata keywords matching “metagen*”, “microbi*”, “bacteria”, or “archaea” was selected for download. Raw sequence data was batch downloaded using the NCBI SRA Toolkit^69^ (v2.8^70^ totalling over 707 terabytes of raw data from 2,390,937 sequencing samples.

### Sample metadata extraction and processing

The NCBI SRA provides biological metadata for samples, but quality and detail of annotations vary across studies since they depend on data submitters. To standardize this metadata, keywords were extracted from multiple annotation fields (isolation source, disease status, host information, body habitat/site, environmental package, and tissue annotations) and processed through string normalization and filtering steps, including lowercasing, removal of special characters, and tokenization. In addition, custom rules for term disambiguation, plural forms and synonyms were implemented and applied. The resulting controlled vocabulary was used to classify samples into four main environments: animal, plant, aquatic, and soil. Samples were assigned to main environments by matching their standardized keywords against environment-specific term sets (e.g., “leaf, banana, tree, crop” matches only “plant”; automatic assignment: “plant”) (Supplementary Table 7). When samples had annotations matching multiple environments, they were labelled as “unknown” (e.g., “leaf, banana, tree, insect” matches both “animal” and “plant”; automatic assignment: “unknown”).

Host species were initially assigned by matching relevant keywords and, for more quantitative analyses, through explicit parsing of dedicated metadata fields containing host names and taxonomic IDs. To this end, relevant metadata fields were detected using regular expressions matching variations of “host”, “name” and “taxid”. Identified fields were manually checked and added to a white list, accounting for misspelling, word separation and capitalization. Entries from white-listed fields for all samples were then parsed and homogenized by translating identified names to NCBI taxonomic IDs via the ete3 toolkit^71^ v3.1. Samples with multiple conflicting host assignments were treated as unassigned.

### OTU clustering and taxonomic classification

The full-length SSU rRNA reference database MAPref 2.2b, delivered with MAPseq 1.2.6^72^, was used to define and taxonomically classify OTUs. Briefly, the database contained 1,536,850 quality-filtered full-length 16S and 18S sequences, extracted from the NCBI RefSeq and Genbank databases. Sequences were aligned and hierarchically clustered using HPC-CLUST^73^. Clustering was performed at different sequence identity thresholds (90%, 96%, 97%, 98%, and 99%), resulting in hierarchical OTU definitions (parents and children). Taxonomies for each OTU were assigned based on a 90% consensus annotation across all of its sequences, obtained either directly via NCBI RefSeq and the All-Species Living Tree Project taxonomic annotations (“gold set”) or indirectly by mapping un-annotated sequences to the gold set (“silver set”). For details, see ^72^.

### Raw read quality filtering and OTU assignment

Fastq sequencing data was extracted from downloaded SRA sequencing runs using the fastq-dump provided by NCBI. A custom C++ program was used to filter and trim the reads using the quality information, based on the following criteria: base calls were defined as low quality when they had a quality score below 10, reads were trimmed when two consecutive low quality base calls were identified, and reads shorter than 75bp or with a fraction of low quality base calls higher than 5% were discarded.

MAPseq v1.2.6 was used to assign all quality-filtered reads to the NCBI taxonomy and to OTU labels at different identity cutoffs (90%, 96%, 97%, 98%, and 99%). A confidence cutoff of 0.5 was used to obtain confident read assignments; reads with alignment scores lower than 30 were discarded.

Samples with less than 20 OTUs (OTU level: 96%), with less than 1000 SSU reads, or not annotated as “source=METAGENOMIC” were discarded to reduce the fraction of lower-quality or non-microbiome samples, resulting in 1,153,349 remaining samples. This dataset served as the base for community clustering (see below). For per-sample downstream analyses, samples with less than 90% estimated community coverage^74^ (based on 97% OTUs) were additionally removed, yielding a dataset of 1,039,362 samples.

### Community clustering

For all samples with at least 20 OTUs and 1000 SSU reads, pairwise microbial community distances were calculated (Bray-Curtis dissimilarity on log-transformed relative abundances), followed by average linkage hierarchical clustering (distance threshold: 0.5). To handle the computational demand of >1 million samples, a modified version of HPC-CLUST was used. This yielded 356,182 community clusters, which were further reduced to a more robust and computationally tractable subset of 36,358 clusters with at least five samples each. A majority rule was used to assign projects, environments, sequencing strategies, hosts IDs, and metadata keywords to community clusters, where terms found in at least 80% of the samples were assigned to the whole cluster. Clusters assigned to project ID ERP012803 were annotated as American Gut Project (AGP), clusters assigned to projects SRP002395, SRP002860, SRP002163, or SRP002012 were annotated as Human Microbiome Project (HMP), and clusters assigned to any project from https://ebi-metagenomics.github.io/blog/2019/04/17/Earth-Microbiome-Project/ were annotated as Earth Microbiome Project (EMP). 26,487 community clusters with confidently assigned main environments and sequencing technologies underwent UMAP projection and follow-up analyses (see subsection “Statistics, normalization, and visualization”).

For environmental annotations in Fig. 2a, our 65 keyword-based sub-environments were manually summarized into up to three categories per main environment. For the analysis of human exposed body sites and human gut microbiomes, community clusters assigned to “Animal (human)” were further annotated as exposed body sites if their consensus keywords included “vagina”, “oral”, “skin”, “saliva”, “salivary”, “human-oral”, “urogenital”, “mouth”, “nare”, “nasal”, “anterior nare”, “lung”, “airways”, or “nasopharynx”. They were instead annotated as gut if their consensus keywords included “gastrointestinal”, “feces”, “gut”, “gastric”, “stomach”, “gastrointestinal tract”, or “human-gut”. To reduce noise, human and environmental clusters in Supplementary Fig. 5 were further refined by applying coordinate masks to their core regions within the projection.

For the mouse and human gut comparisons, only clusters annotated as “Animal (human)” or “Animal (mouse)” were considered. Core cluster boundaries were refined via hierarchical density clustering with HDBSCAN^75^ v0.8.40. Keywords enriched in human and mouse, respectively, were computed using Fisher’s exact test (two-tailed) to remove noisy and uninformative terms. Significantly enriched and depleted keywords were then pooled and semantically clustered. To this end, the large language model (LLM) Llama-3.2-3B-Instruct^76^ was iteratively provided with sets of 20 randomly chosen keywords and then asked to group these keywords into clusters of semantically related terms. This involved i) a system prompt that confined the LLM’s response to an easily parsable format and ii) a user prompt with the clustering instructions. Through this repeated process, we sampled the LLM’s implicit relational knowledge, allowing for the creation of dataset-specific keyword semantic distances. Next, the semantic distance for each pair of keywords was computed as:

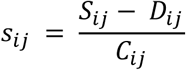

Where *s* is semantic distance between keywords *i* and *j*, *S* represents the number of contexts in which *i* and *j* cluster together, *D* represents the number of contexts in which they cluster apart, and *C* represents the total number of contexts in which they both appear. Intuitively, keywords that are more closely related within the set of possible keywords should cluster together more often on average, decreasing semantic distance.

Based on these semantic distances, umap^77^ (v0.5.7) and sklearn^78^ (v1.5.2) K-means clustering were used to reduce and identify clusters of semantically related keywords. For each cluster, the average odds ratio, defined as average fold change of prevalence in mouse compared to human samples, was calculated.

### Geographic mapping and IUCN annotation

Sampling coordinates (latitude and longitude) were parsed directly from dedicated SRA annotation fields; a total of 31 distinct field headers were encountered that either matched ‘longit*’ or ‘latitud*’ separately, or a combination of ‘lat*’ and ‘lon*’. For the coordinates themselves, a variety of different formatting choices were observed, such as decimal vs. degree notations, various levels of precision, different syntaxes, and different ways of distinguishing North/South and East/West. The coordinates were thus parsed using distinct regular expressions in custom Python code; in an iterative procedure, coordinate records that initially failed to parse were inspected manually and corresponding regular expressions for special cases were added (a total of 11 regular expressions were eventually needed).

In a subsequent step, the resulting sample placements were inspected visually on a zoomable world-map; large sample collections with improbable coordinates stood out and were followed up manually. Obvious errors were corrected if possible; these included accidental lat/lon reversals in the original metadata, as well as series of samples that differed systematically by one numerical unit from each other, creating vertical or horizontal lines of samples on the map (these possibly originate from MS Excel manipulation errors).

For IUCN Global Ecosystem Typology assignments, the filtered MicrobeAtlas dataset was further screened for samples assigned as “soil” and with available coordinates, yielding 98,642 samples. Indicative geographic maps for terrestrial Ecosystem Functional Groups (EFGs) were obtained from the IUCN^44^ (https://global-ecosystems.org/). Maps with lower spatial resolution (T2.5 “Temperate pyric humid forests” and T1.3 “Montane tropical forests”) were improved by integrating more recent maps^79,80^ for EFG-specific criteria applied by the IUCN, yielding resolutions of 30S per pixel. Samples were only assigned to an EFG if i) their coordinates matched an area of “major occurrences” (covering >=25% surface area) of that EFG, and ii) no other EFGs had a “major occurrence” in that area. Similarly, terrestrial IUCN Biomes were assigned if a sample’s coordinates overlapped with “major occurrence” EFGs from only a single biome.

### Cultured isolate and genome mapping

Two external databases were crosslinked to MicrobeAtlas: proGenomes3^46^ (accessed: September 2022), containing almost one million bacterial genomes, and BacDive^45^ (accessed: November 2022), containing genomes of bacterial strains in culture. To estimate how much of the diversity contained within MicrobeAtlas is covered by these two resources, 16S rRNA genes were extracted from the genomes of both proGenomes3 and BacDive. When multiple 16S rRNA genes were found in proGenomes3, the longest sequence was selected (with a minimum length of 600 bp). Out of 907,388 genomes in proGenomes3, 753,909 representative 16S rRNA sequences were extracted using Barrnap^81^. These were then mapped together with 16S sequences from BacDive (based on annotations within NCBI GenBank and RefSeq) to the 97%-identity set of MicrobeAtlas OTUs. Mapping was performed with MAPseq against the MAPref 3.0 reference set, keeping only best hits with a minimum identity of 97%. Following this procedure, 732,912 of the 753,604 (97.3%) 16S rRNA sequences identified in proGenomes3 confidently mapped to 11,084 MicrobeAtlas 97%-OTUs. In BacDive, 26,269 of the 28,227 (92.6%) sequences were confidently mapped to 6,722 MicrobeAtlas 97%-OTUs.

The extracted 16S sequences from proGenomes3 underwent further analysis to establish links between specI clusters and MicrobeAtlas OTUs. The linking was performed at the 98% OTU level (approximately species level), which in our tests optimized the overlap between specI clusters and MicrobeAtlas OTUs. A majority-rule approach was implemented, where a connection was only established when a minimum of 80% of the 16S sequences (mapped with at least a 40% confidence score) within one specI cluster corresponded to the same 98% MicrobeAtlas OTU. This process resulted in 19,902 specI clusters reliably mapped to 9,511 MicrobeAtlas 98%-OTUs.

To ensure a finer sequence resolution and to better reflect the strain level differences contained in BacDive, 99%-OTUs were used for the MicrobeAtlas-BacDive links reported on the website (https://microbeatlas.org). Only unambiguous mappings with at least a 40% confidence score were kept, resulting in 15,656 BacDive strains linked to 9,434 MicrobeAtlas OTUs at the 99% OTU level.

For genome size analyses, MAPref 3.0 OTUs were mapped back to MAPref 2.2b OTUs to ensure compatibility with other analyses in this study. To achieve this, OTUs with high sequence overlap between the two references were linked, resulting in 7427 MAPref 2.2b OTUs with at least one mapped non-MAG genome and associated generalism scores. For each genome, genes were called and counted by running Prodigal^82^ (v2.6.3) with the following parameters: translation table 11 (-g 11), closed ends (-c), treat runs of N as masked sequence (-m), single procedure (-p single).

### Diversity estimation and latitudinal analyses

Total community richness was extrapolated using formula 7 in ^74^ (based on an improved version of the Good-Turing frequency estimator). For additional consistency checks, sample richness was also estimated by counting observed OTUs in samples rarefied to 1000 SSU reads. Beta diversity between samples and community clusters was computed as Bray-Curtis distance on relative abundances, using the Distances.jl package^83^ (v0.10). Weighted and unweighted unifrac dissimilarities were computed using Strided State Unifrac^84^ from the biocore.unifrac package (v1.3). For community clusters, beta diversity between two clusters A and B was defined as the average pairwise Bray-Curtis or unifrac dissimilarity between all samples in A and all samples in B. For unifrac, each community cluster was reduced to three randomly chosen samples to improve computational tractability. The phylogenetic tree used for unifrac was computed by aligning all 16S reference sequences with Infernal (for details, see ^72^), followed by maximum-likelihood tree inference using FastTree^85^ v2.1.10 (parameters: “-nt -gtr -gamma”).

For the latitudinal diversity analysis of marine communities, only samples with sub-environment “marine”, “ocean”, or “sea” were considered (50,480 samples). The “Global oceans and seas” shapefile^86^ (version 1) was used to remove samples not located in oceans, resulting in a final set of 43,156 samples. For latitudinal diversity analysis of terrestrial samples, diversity curves were computed per IUCN biome in order to not mix vastly different ecosystems (e.g., agricultural soils and deserts) within one analysis. Only biomes covering at least 3 of 10 latitudinal bins with a minimum of 100 samples per bin were included. These criteria were only matched by “Deserts and semi-deserts” (600 samples, 3 bins), “Temperate-boreal forests and woodlands” (2,464 samples, 4 bins), and “Intensive land use” (42,133 samples, 7 bins).

To compute latitudinal abundance associations, latitudes were discretized into 50 evenly sized bins and 97% OTUs present in at least 25 bins and with minimum global prevalence of 250 were considered. For each OTU, relative abundances per bin were averaged and log10-transformed, followed by computing Pearson’s ρ between log-transformed averages and per-bin latitudes. OTUs were assigned to a main environment based on their maximum average abundance within the four major environments: animal, aquatic, soil, and plant.

### Habitat generalism

To obtain habitat generalism scores for all OTUs, average relative abundances in each main environment (animal, aquatic, soil, plant) were computed. While computing abundances independently per environment reduces the impact of sampling bias, the choice of relative abundance instead of prevalence reduces the effect of contaminants, which are expected to disperse widely, but to only reach low abundances in non-native samples. These per-OTU environmental abundance profiles were then converted into proportions to make profiles of abundant and rare OTUs comparable. Finally, Shannon entropy scores were computed for each normalized profile, resulting in a measure that peaks if an OTU is found in all major environments with equal abundance (highly generalist) and becomes zero when an OTU is only found in one major environment (highly specialist). OTUs that peak in a few environments but show low or zero abundance in others yield intermediate scores. Scores were further normalized to lie between 0.0 and 1.0 through division by the theoretical maximum entropy (obtained for a perfectly even distribution across the four major environments). Per-OTU main environment assignments followed the same approach as described in the latitudinal analyses section.

### Statistics, normalization, and visualization

UMAP projections were computed based on the Bray-Curtis distances used for community clustering (see above), utilizing the umap package^77^ (v0.5) in julia^87^ v1.9 via PyCall.jl^88^. Parameters were as follows: n_neighbors=500, min_dist=3, spread=35, n_epochs=2500.

Plots were generated using matplotlib^89^ v3.7, seaborn^90^ v0.13, and VegaLite.jl^91^ v2.6. Community clusters were further density-clustered for illustration using DBSCAN, implemented in Clustering.jl^92^ v0.15, with default parameters. OTU dendrograms were generated using the ete3 toolkit^71^ v3.1 via PyCall.jl. Word clouds were computed and visualized using the wordcloud package^93^ (v1.9) in Python and geographic maps were generated on https://microbeatlas.org or in Python using folium^94^ v0.15.1.

Statistical tests and effect sizes throughout the manuscript were computed using HypothesisTests.jl^95^ v0.11 and scipy^96^ v1.10. Where appropriate, Benjamini-Hochberg correction^97^ was used to adjust p-values for multiple testing (significance level: 0.05). Permutational multivariate analysis of variance (PERMANOVA) results were computed with the permanova function from the scikit-bio package^98^. Locally weighted scatterplot smoothing (LOWESS) curves were computed using the “lowess” option in the Python seaborn package, linear regressions and confidence intervals were fitted using the regplot function in seaborn with default parameters.

Unless otherwise stated, relative OTU abundances were used for analyses: per-sample OTU abundances were normalized to a range of 0 to 1 by dividing by the total number of mapped reads. Logarithms throughout the paper were computed by adding a pseudo-count equal to the minimum non-zero value divided by 10 to all values.

## Acknowledgements

We thank all present and former members of the von Mering lab, as well as Maarten Langen, Hans-Joachim Ruscheweyh, and Shinichi Sunagawa for shared data and helpful discussions. We furthermore thank members of the Bork group at EMBL Heidelberg (Chan Yeong Kim, Daniel Podlesny, Jonas Schiller, Peer Bork) for critical reading of the manuscript before submission. The work was funded by the Swiss National Science Foundation (project grant 310030_192569; the Swiss National Centre of Competence in Research (NCCR) Microbiomes, 310030_192567).

## Author contributions

Conceptualization: J.F.R., J.T., C.v.M.; Data download and processing: J.F.R., J.T., L.M., D.P., E.P.M.M., T.S.S., C.v.M.; Scientific and bioinformatic analysis: J.T., J.F.R., L.M., E.P.M.M., N.N., M.E.P., Ma.D.; Website: J.F.R., D.P., G.R., Mi.D., C.v.M.; Figure creation: J.T., J.F.R., L.M., E.P.M.M., N.N., D.G., C.v.M.; Supervision: J.T., J.F.R., C.v.M.; Manuscript writing: J.T., J.F.R., C.v.M.; Manuscript feedback: all authors.

## Data and code availability

This paper analyzes existing, publicly available data, accessible at www.ncbi.nlm.nih.gov/sra. Primary processed data is available at www.microbeatlas.org/download. Original code for downstream analyses has been deposited at https://github.com/meringlab/MicrobeAtlas_compa <u>nion</u> and is publicly available as of the date of publication. Any additional information required to reanalyze the data reported in this paper is available from the lead contact upon request.

## Declaration of generative AI and AI-assisted technologies in the writing process

During the preparation of this work the authors used ChatGPT 4o^99^ in order to improve conciseness and clarity in parts of the main text and legends. After using this tool/service, the authors reviewed and edited the content as needed and take full responsibility for the content of the publication.

## Supplementary figures

**Supplementary Figure 1:**
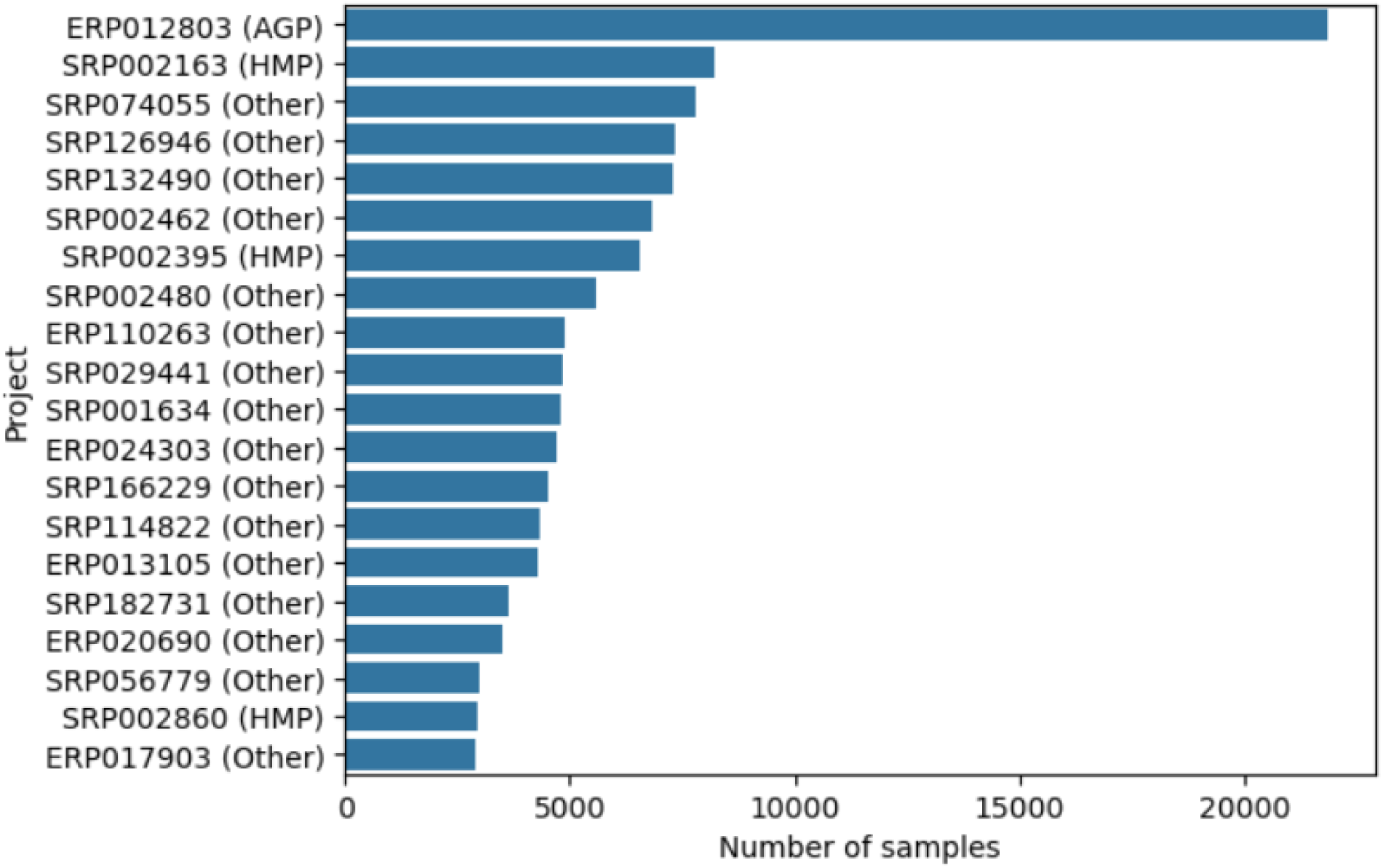
Number of samples in the top 20 projects in MicrobeAtlas. Sample counts are based on a filtered MicrobeAtlas subset of 1,153,349 sequencing samples, excluding those with low complexity or low sequencing depth. AGP: American Gut Project, HMP: Human Microbiome Project, Other: projects not assigned to AGP, HMP, or EMP (Earth Microbiome Project).

**Supplementary Figure 2:**
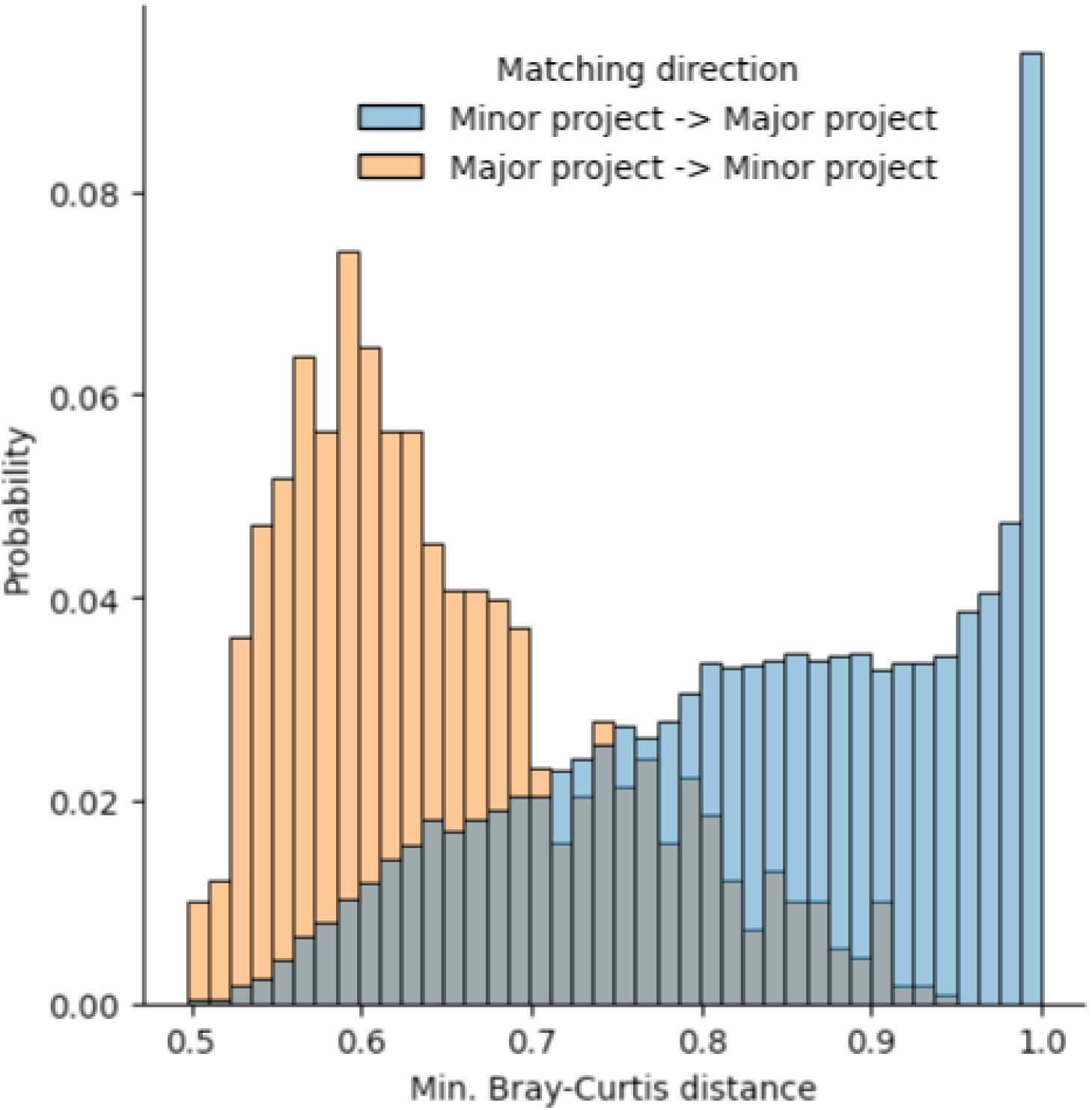
Matching of closest samples between major and minor projects. Bray-Curtis distances between samples from major projects (Human Microbiome Project, Earth Microbiome Project, American Gut Project) and minor projects (all remaining studies), computed at the level of community clusters. Only samples assigned to robust community clusters (min. 5 samples per cluster) and that did not include both major and minor project samples were considered. For each major project sample, we computed the Bray-Curtis distance between its community cluster and the closest minor project community cluster (orange histogram). Conversely, for each minor project sample, we determined the distance between its community cluster and the nearest major project community cluster (blue histogram).

**Supplementary Figure 3:**
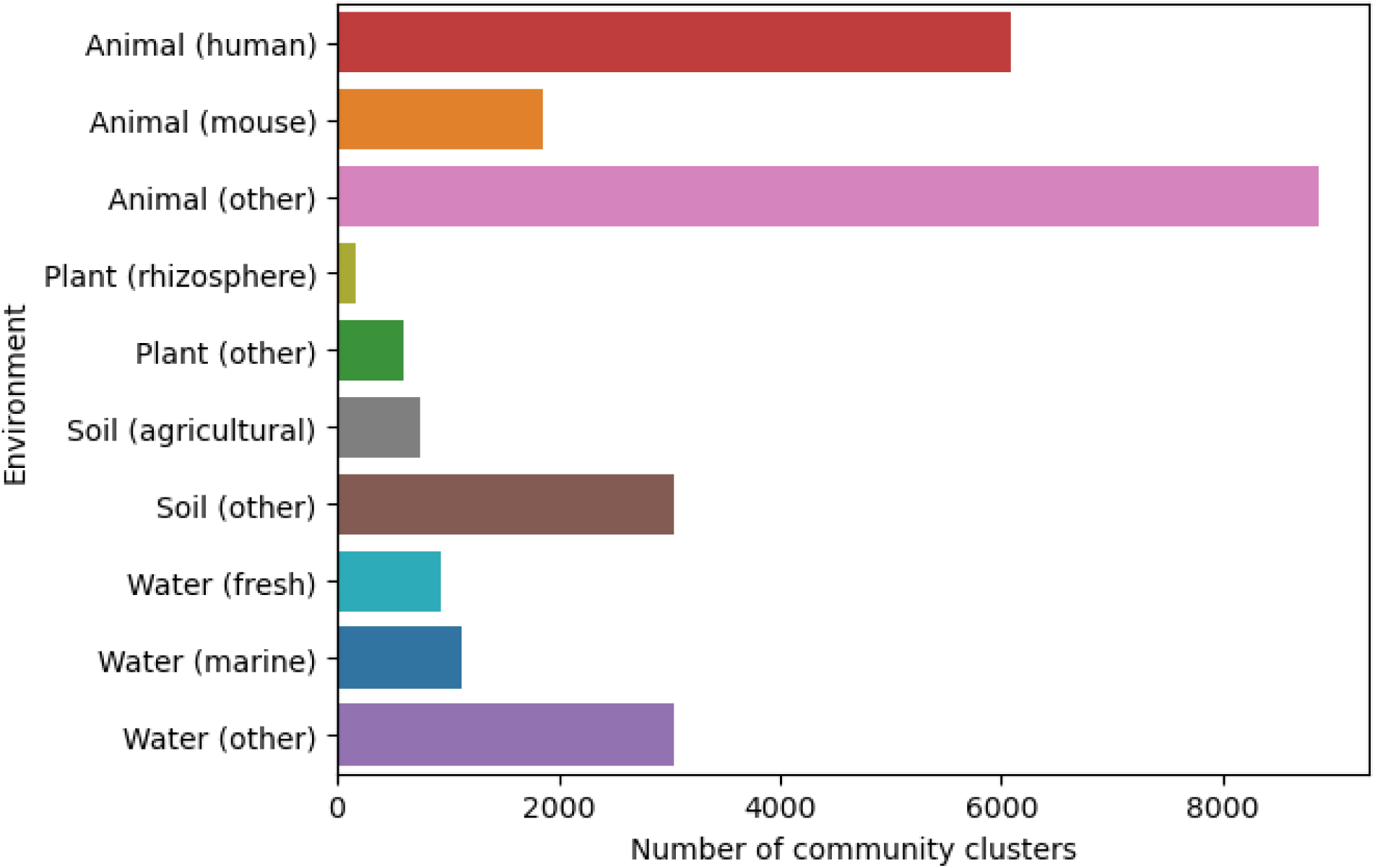
Number of community clusters per environment. “other” includes clusters without sub-environment annotation, clusters with no confident main environment were removed.

**Supplementary Figure 4:**
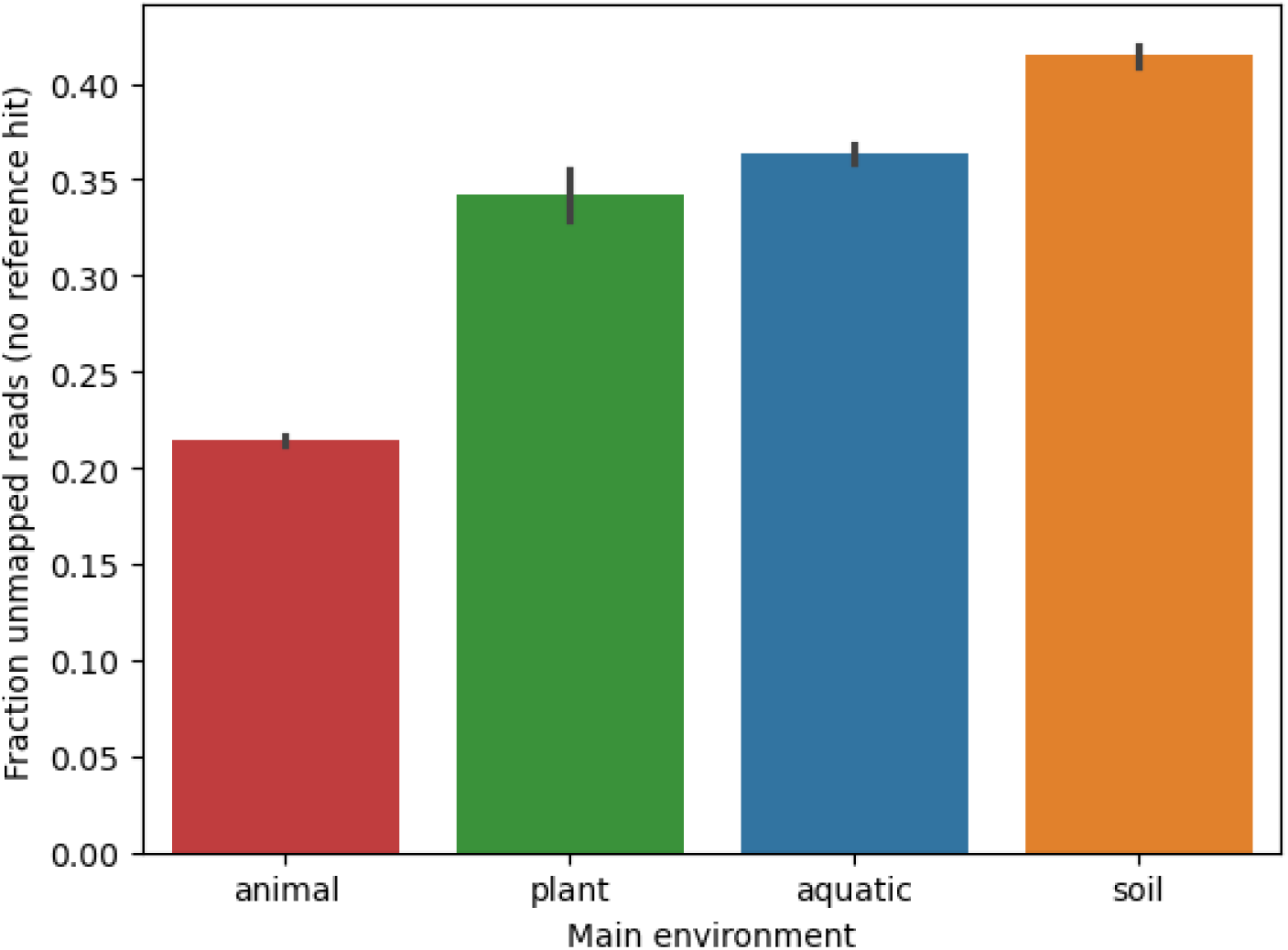
Fraction of SSU reads with no reference hits for each by main environment. Fractions were averaged across robust community clusters (min. 5 samples per cluster).

**Supplementary Figure 5:**
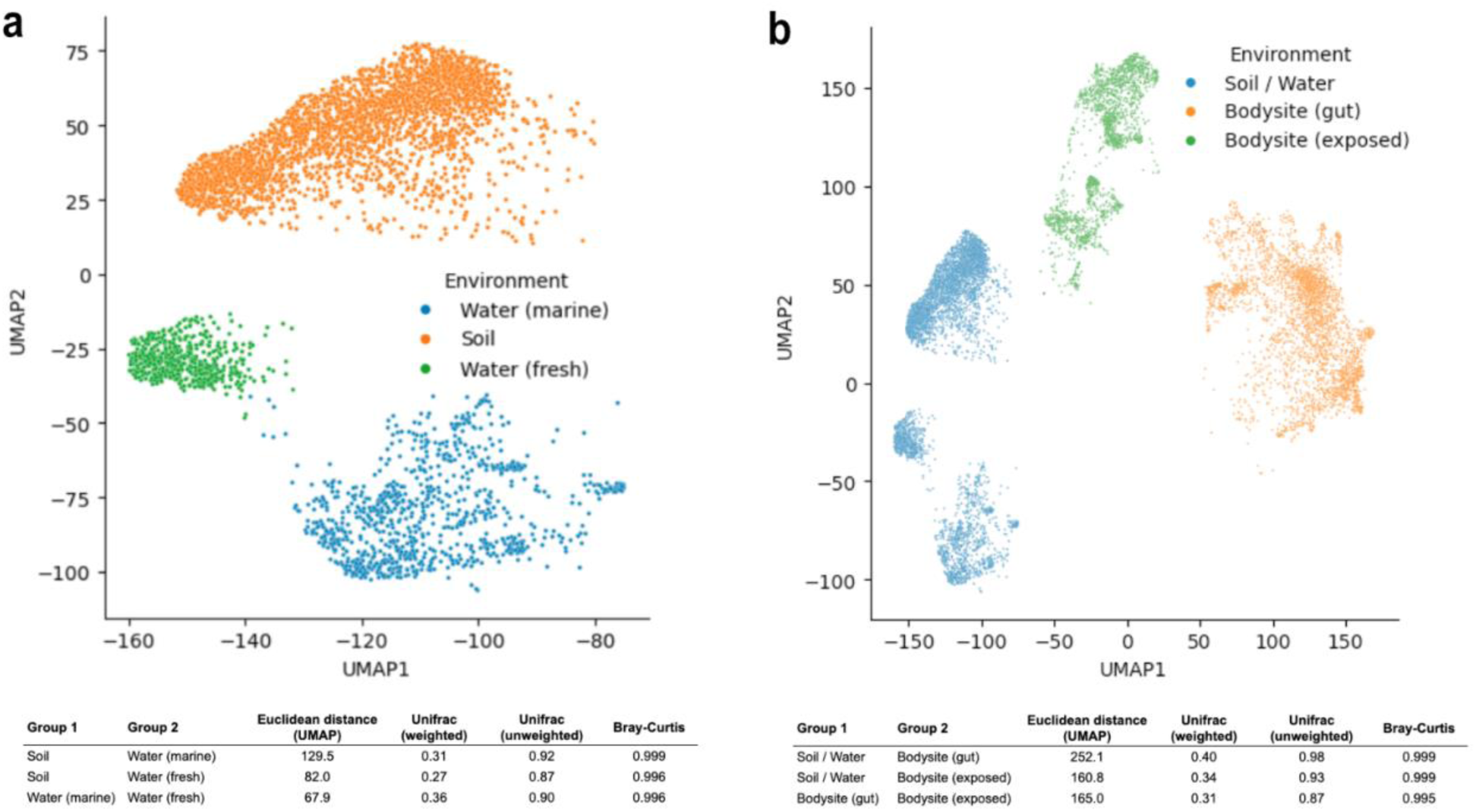
Distances between environmental and body site communities. a, Core clusters of soil, fresh water, and marine communities are shown, together with their average euclidean distances within the projection (bottom table). **b,** Core clusters of more exposed human body sites (skin, oral, nasal/airways, vagina) and human gut are shown in addition to clusters from (a), together with euclidean projection distances between human and environmental clusters (bottom table).

**Supplementary Figure 6:**
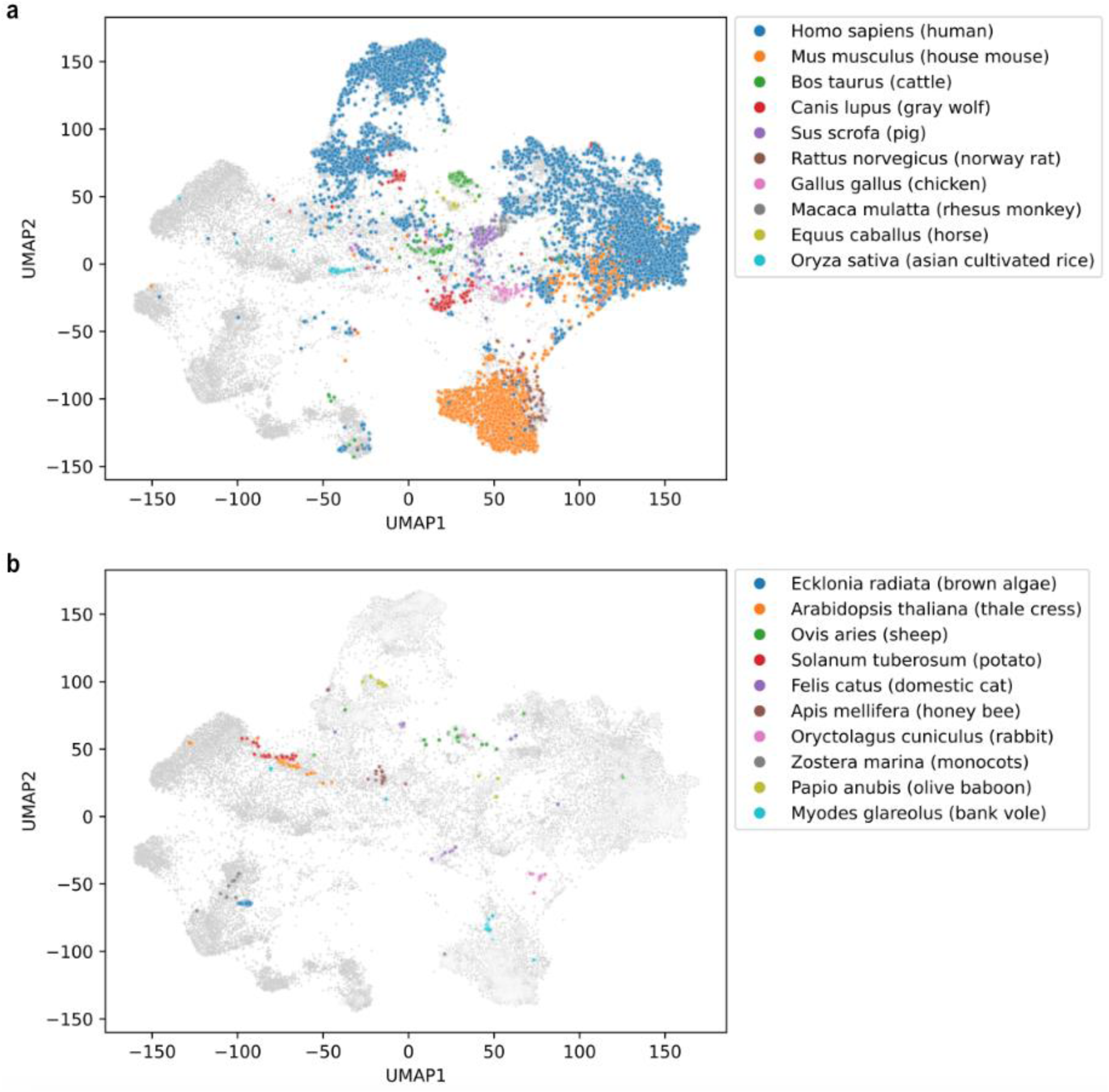
Distribution of host-specific community clusters in a global projection. Community clusters (dots) based on Bray-Curtis dissimilarities, colored by the top 10 (a) and top 11-20 (b) most frequent hosts.

**Supplementary Figure 7:**
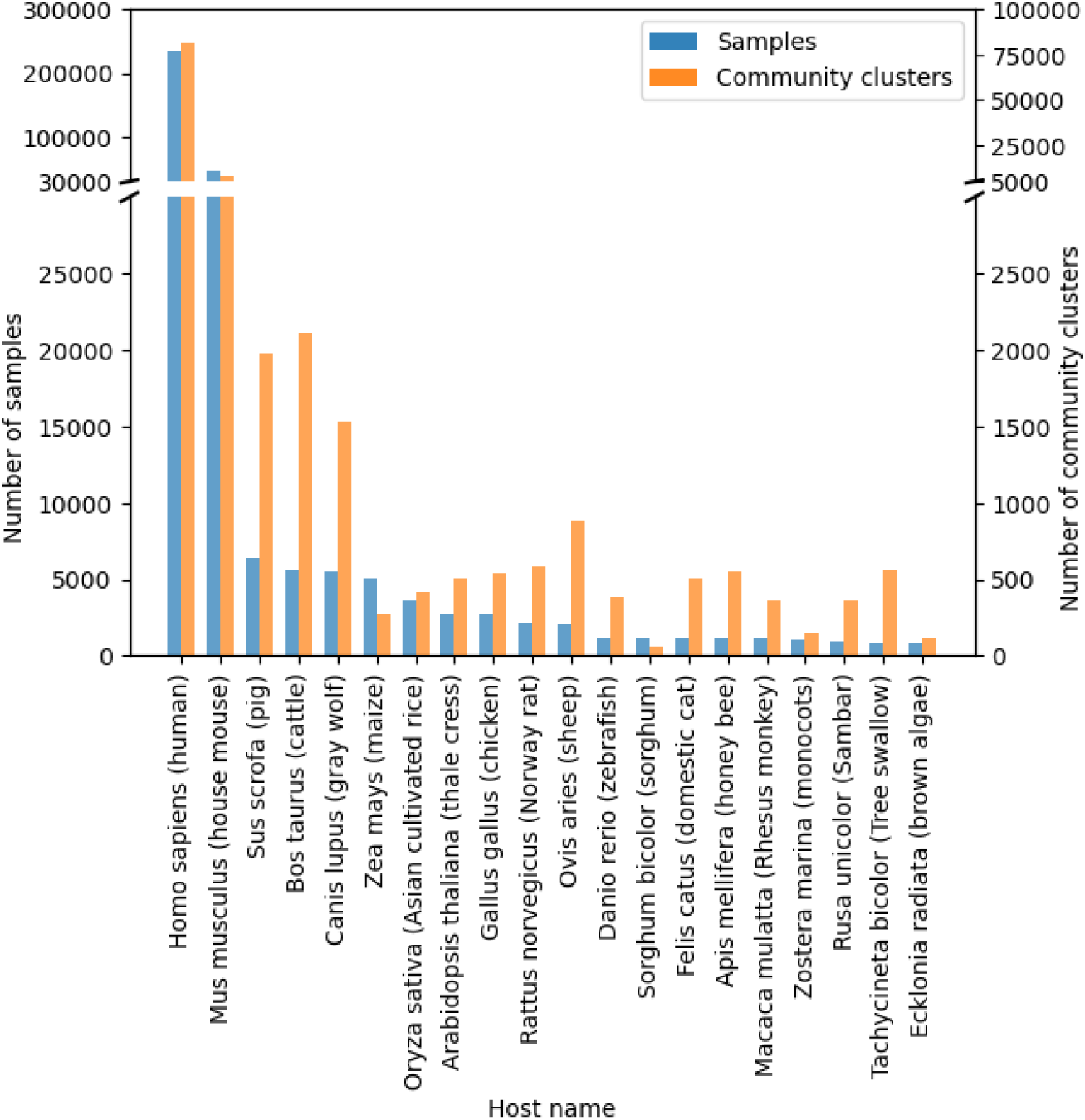
Top 20 most frequently observed host species. Host assignments are based on dedicated metadata fields and taxonomic IDs (Methods). Sample counts are based on a filtered MicrobeAtlas subset with 1,153,349 sequencing samples, excluding those with low complexity or low sequencing depth.

**Supplementary Figure 8:**
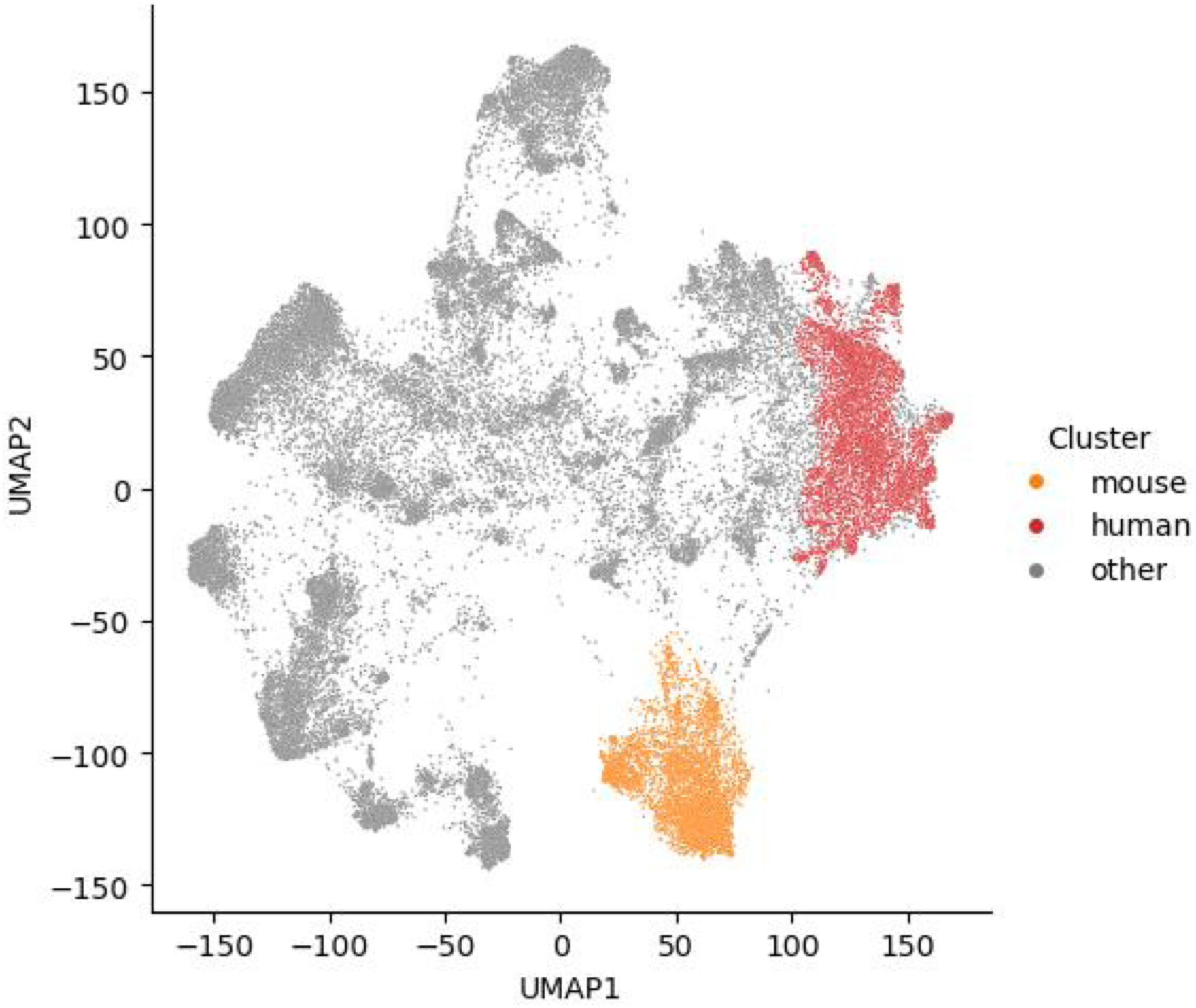
Core mouse gut and human gut density clusters. A parameter sweep to maximise a combined function for human and mouse gut cluster quality based on coverage and purity was performed. With the best performing parameters, the top 5 human gut purity clusters were merged then visualised with the top mouse gut purity cluster.

**Supplementary Figure 9:**
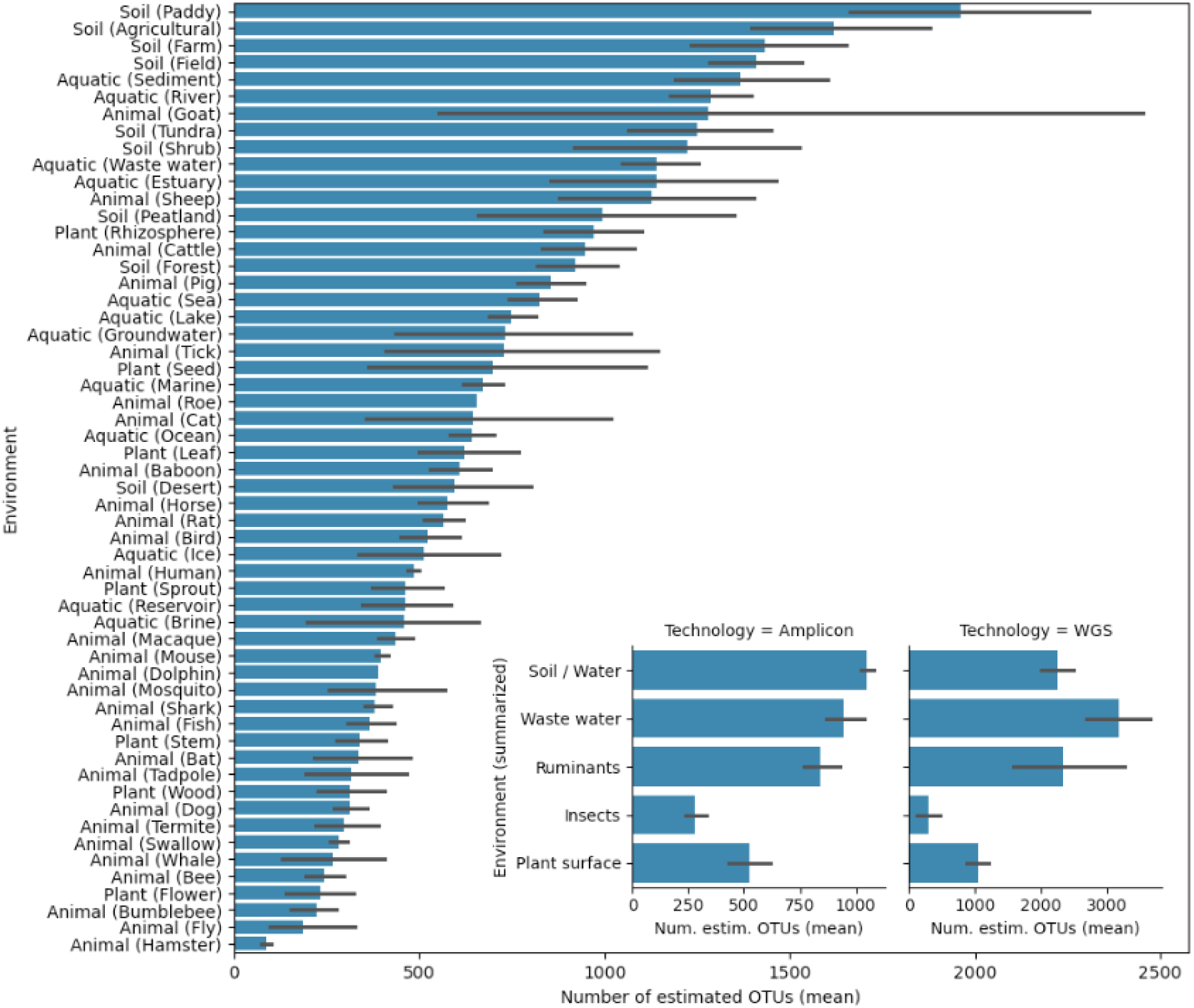
Estimated community cluster richness across all sub-environments. Estimated OTU richness (Methods), averaged per community cluster, for each sub-environment assigned to at least one robust community cluster (min. 5 samples). Inset: richness estimates for manually summarized categories, separately for amplicon and whole genome shotgun (WGS) experiments.

**Supplementary Figure 10:**
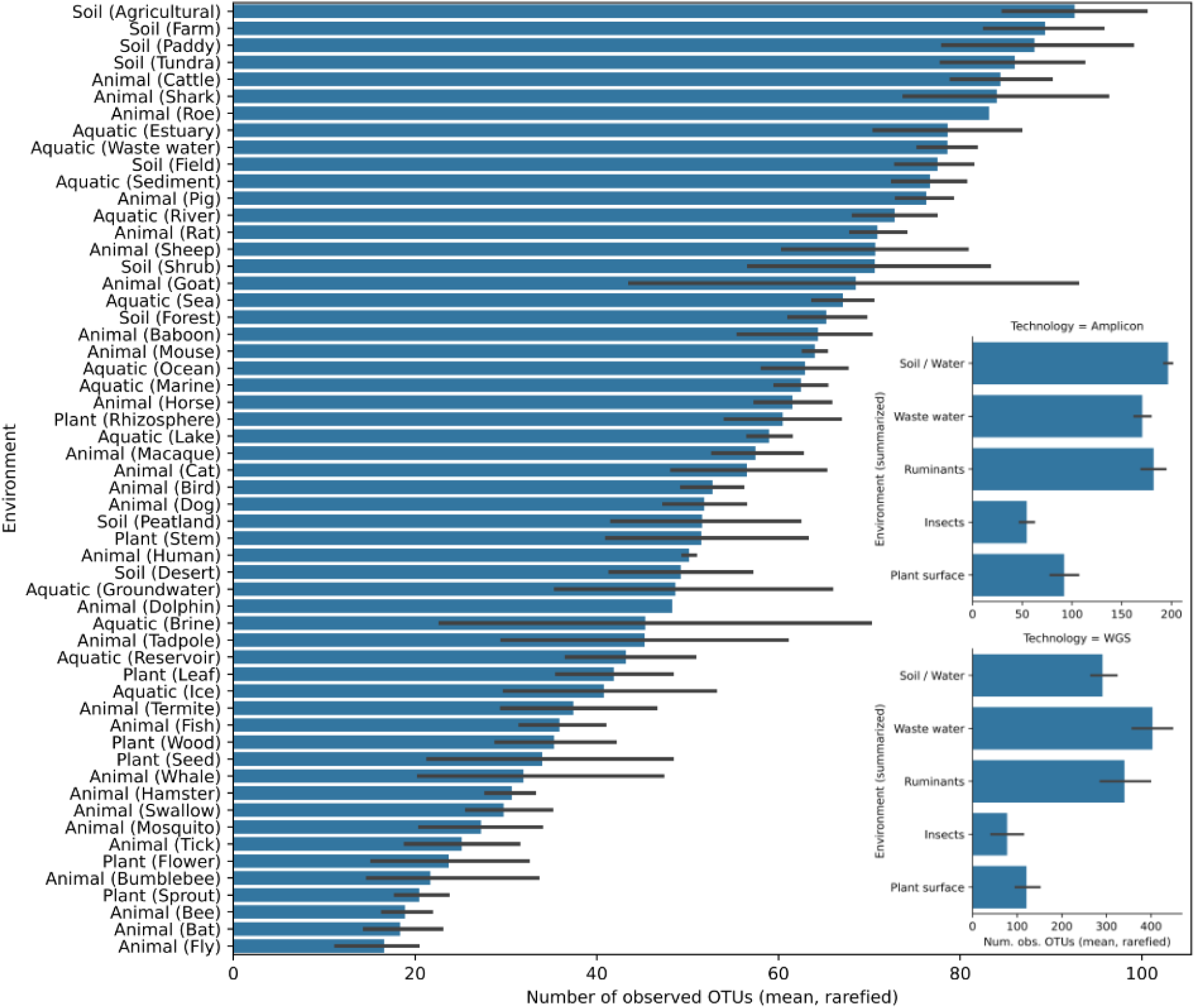
Observed rarefied community cluster richness across all sub-environments. Observed OTU richness (samples uniformly rarefied to 1000 SSU reads), averaged per community cluster, for each sub-environment assigned to at least one robust community cluster (min. 5 samples). Inset: observed richness for manually summarized categories, separately for amplicon and whole genome shotgun (WGS) experiments.

**Supplementary Figure 11:**
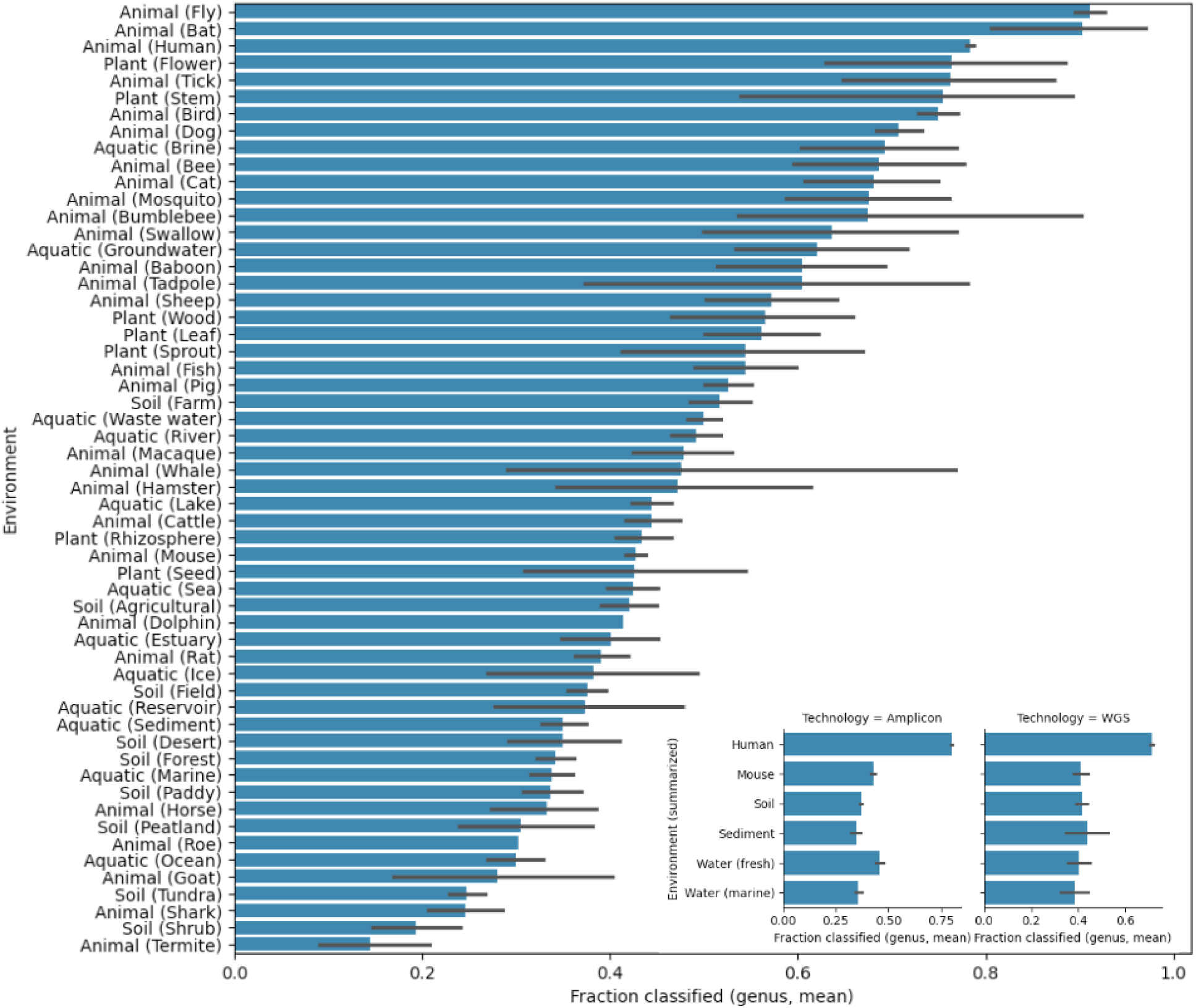
Genus-level characterization of community clusters across sub-environments. Fraction of reads classified at the genus level, averaged per community cluster, for each sub-environment assigned to at least one robust community cluster (min. 5 samples). Inset: genus-level classification fractions for manually summarized categories, separately for amplicon and whole genome shotgun (WGS) experiments.

**Supplementary Figure 12:**
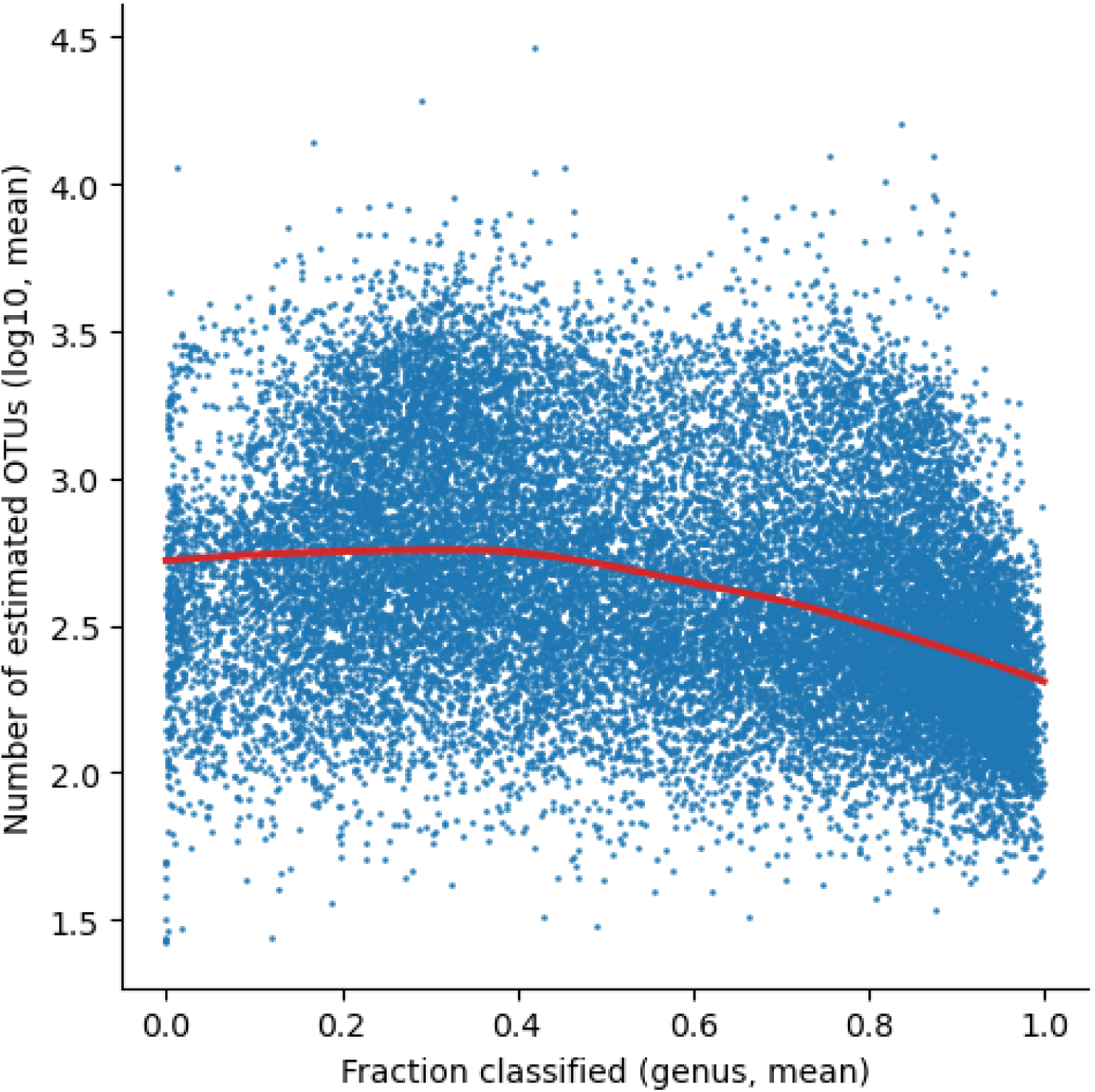
Relationship between richness and classification depth across community clusters. Each dot represents a community cluster, and the curve was fitted using locally weighted scatterplot smoothing (LOWESS), a non-parametric regression method.

**Supplementary Figure 13:**
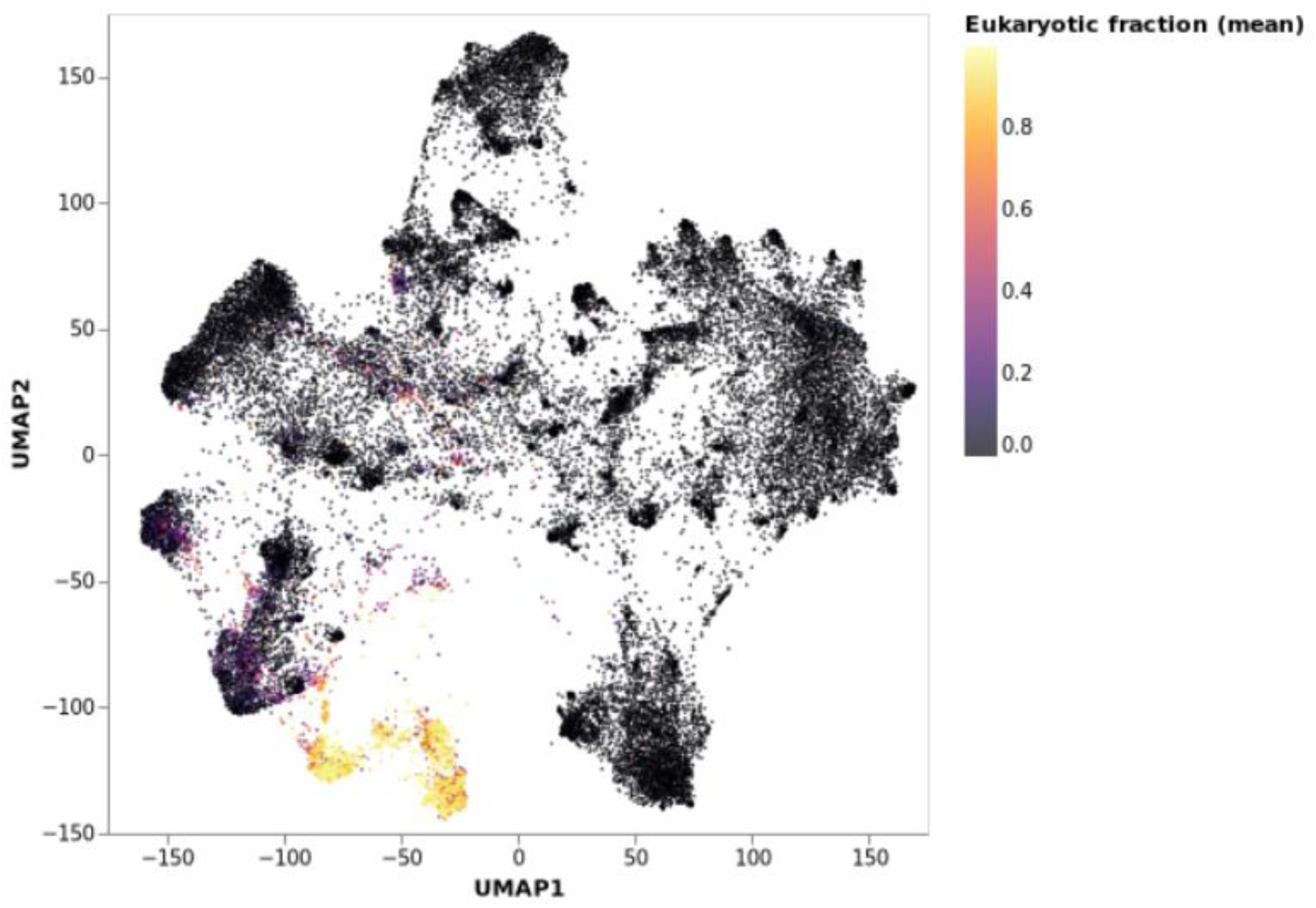
UMAP projection of community clusters colored by the average fraction of eukaryotic OTUs. Eukaryotic fraction was calculated as the average fraction of mapped reads that were assigned to eukaryotic OTUs.

**Supplementary Figure 14:**
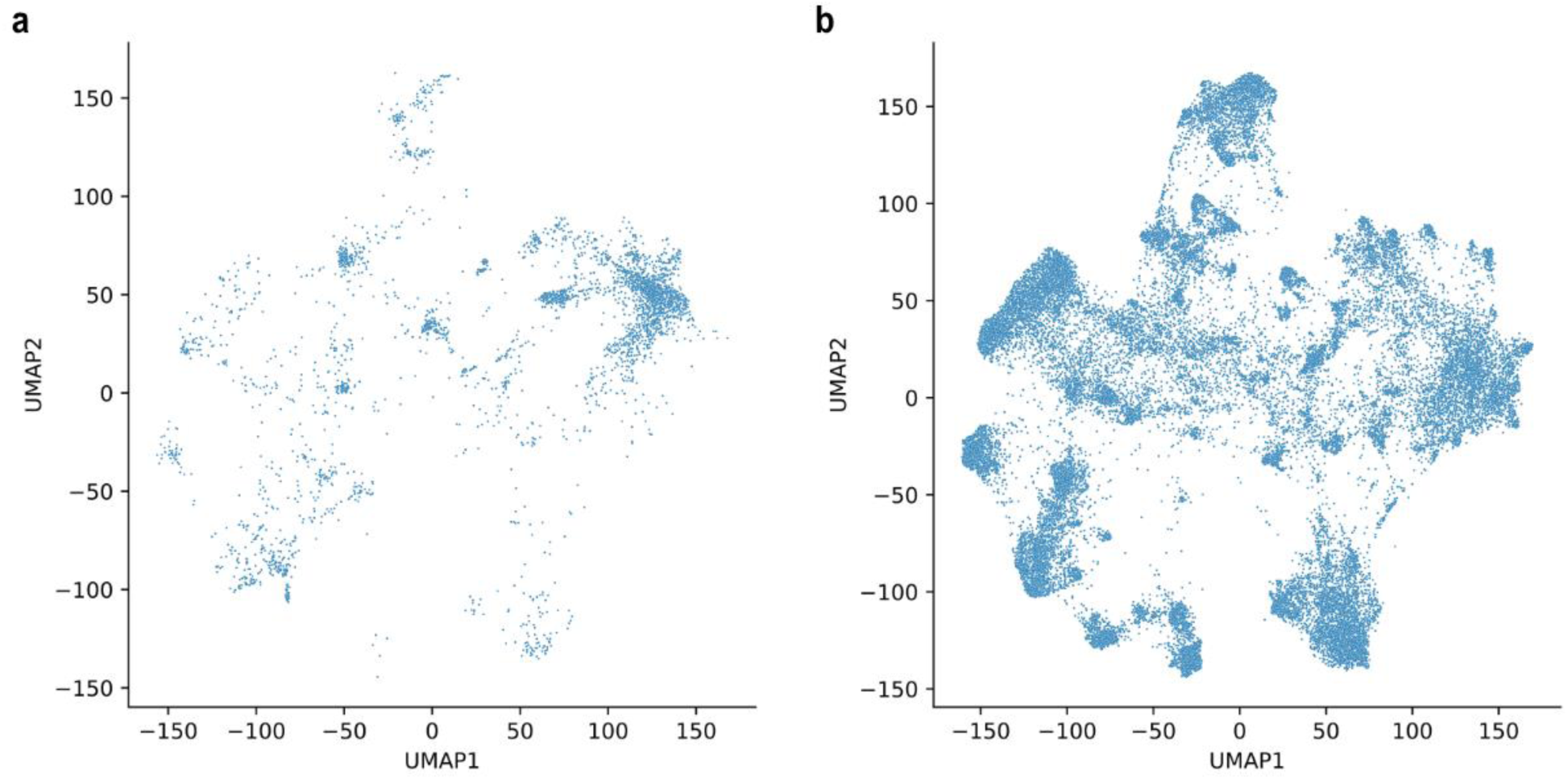
UMAP projection of community clusters by class of sequencing technology. Community clusters generated predominantly (>80% of samples) with whole-genome shotgun (a) or amplicon (b) technologies.

**Supplementary Figure 15:**
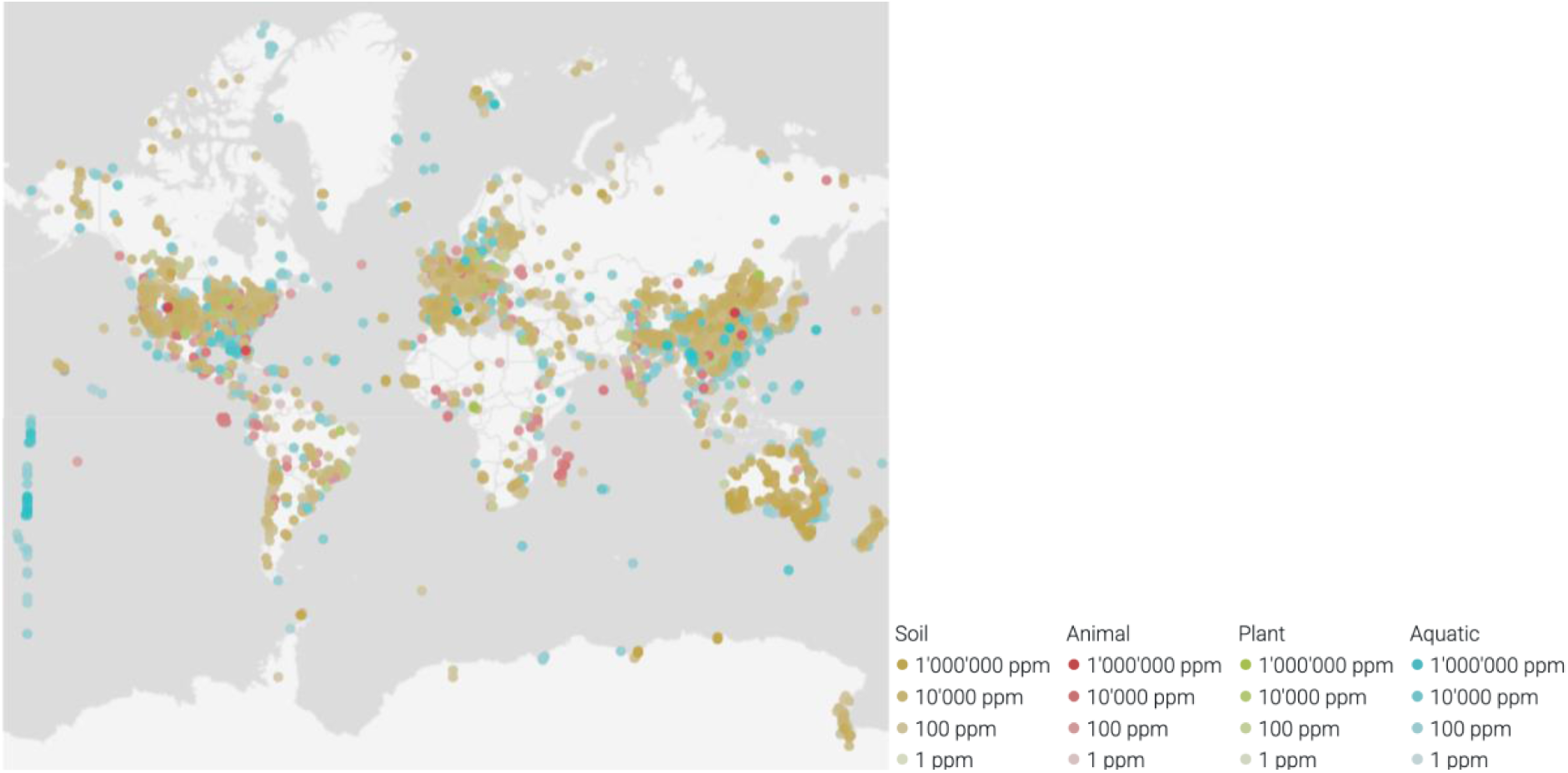
Global abundance distribution of OTU 97_89870 (*Candidatus* Nitrosocosmicus). Abundances shown in parts per million (ppm), with each dot representing an individual sample.

**Supplementary Figure 16:**
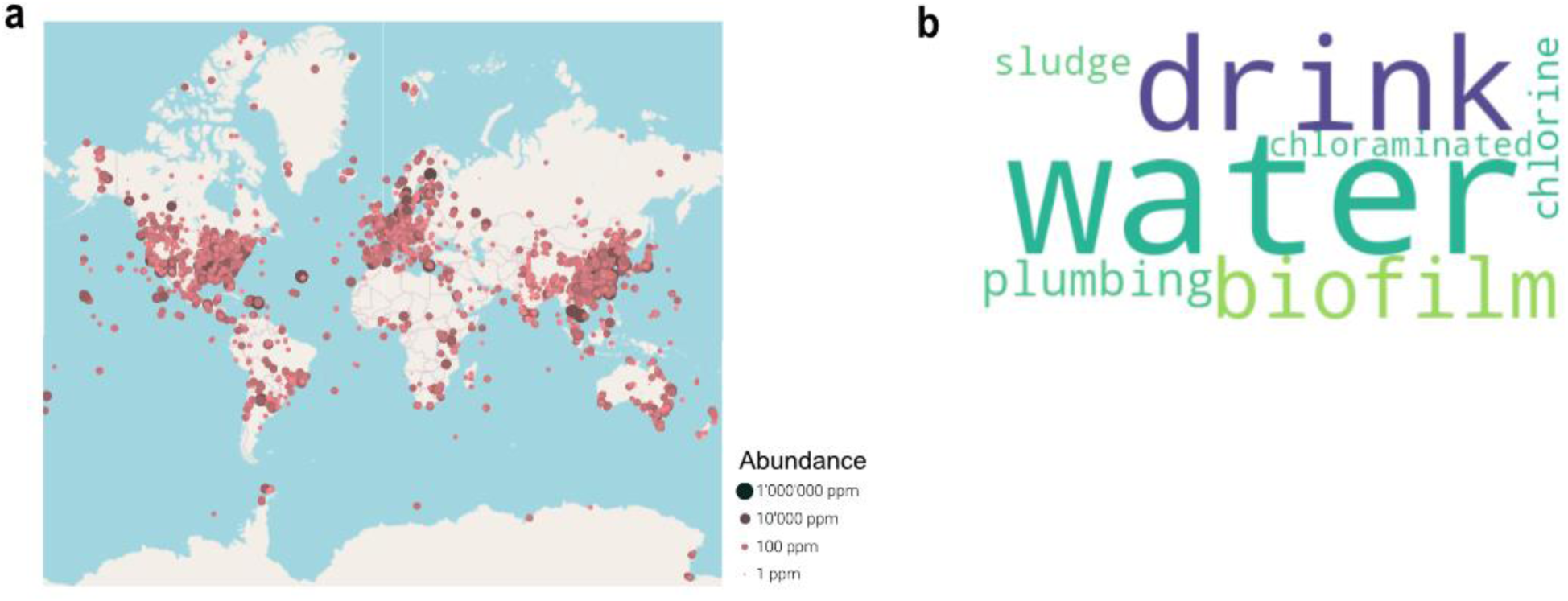
Global distribution of the uncharacterized OTU 97_17665. **a**, Abundance of OTU 97_17665 across geographic regions, shown in parts per million (ppm), with each dot representing an individual sample. **b**, Word cloud of the seven most prevalent keywords for OTU 97_17665 across inhabited samples, weighted by its relative abundance per sample. Text size according to abundance-weighted keyword frequency.

**Supplementary Figure 17:**
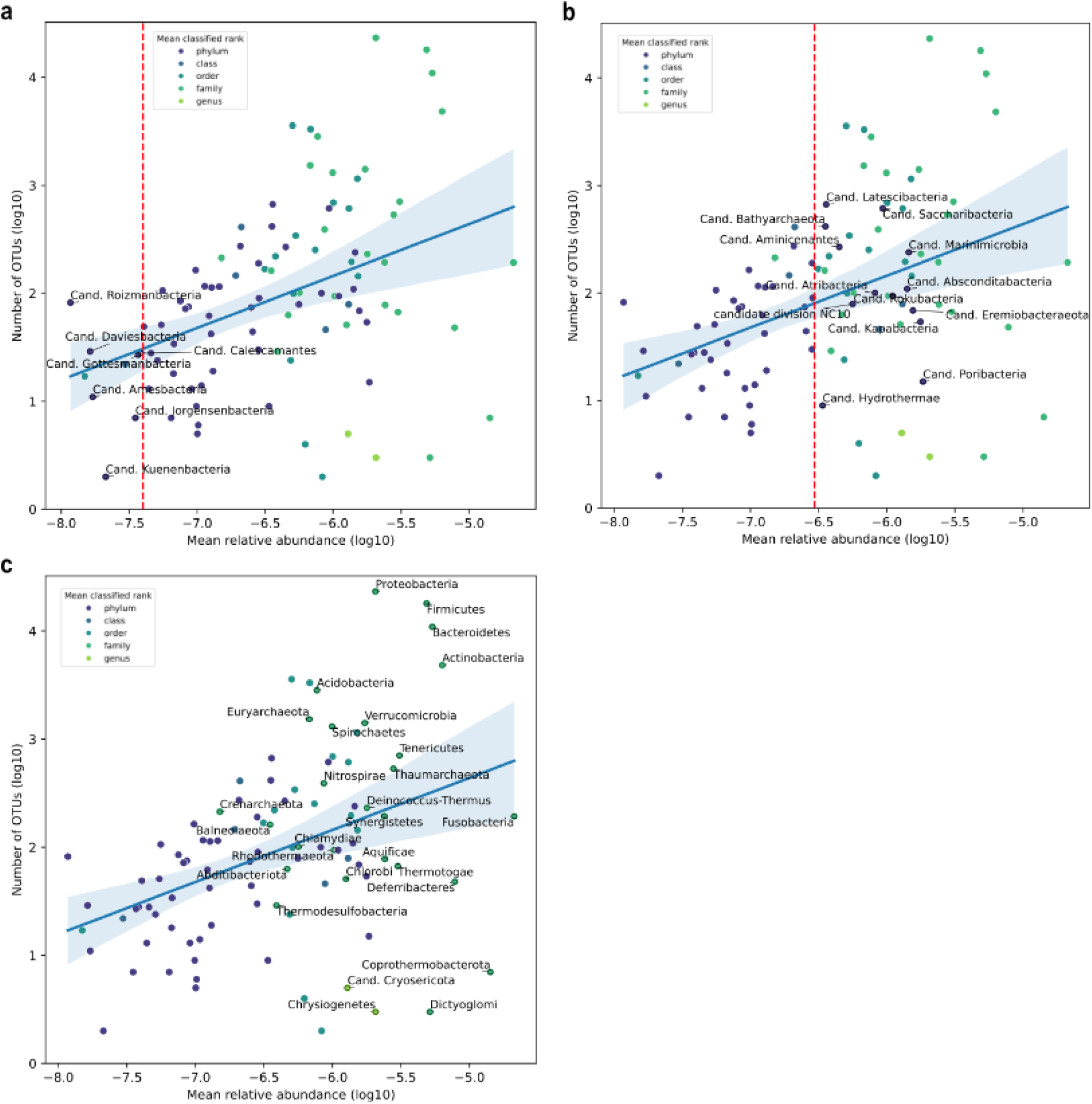
Diversity and global abundance across taxonomic phyla. Each dot represents a phylum. The y-axis shows the number of 97% OTUs assigned to each phylum, while the x-axis represents their mean global abundance. Colors indicate the average taxonomic rank at which OTUs from each phylum are classified. (a) and (b) highlight largely uncharacterized phyla (where OTUs are classified only at the phylum level) with low and high abundance, respectively. (c) highlights better-characterized phyla, where OTUs are classified at the genus or family level. Error bands around linear regression lines represent 95% confidence intervals of log-transformed numbers of OTUs (1000 bootstraps).

**Supplementary Figure 18:**
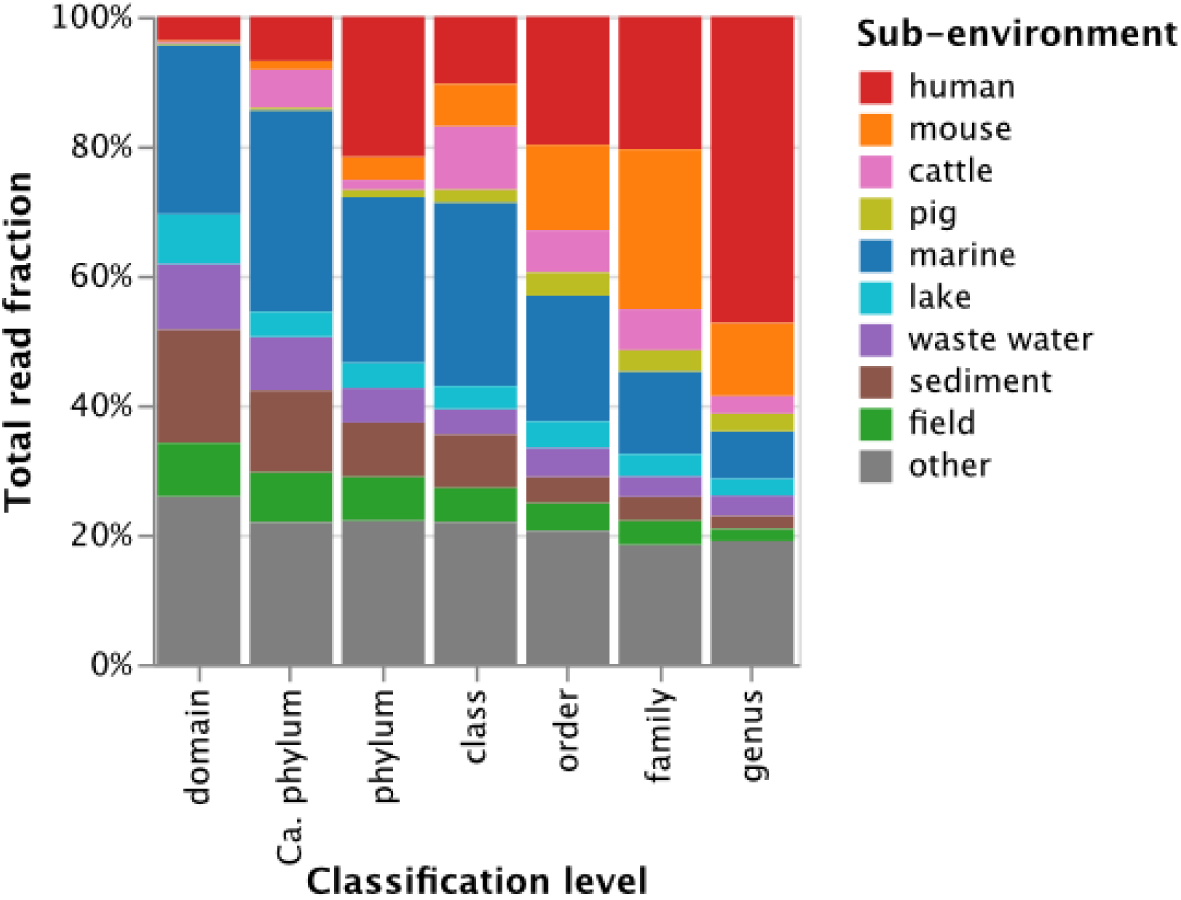
Sub-Environmental distributions of OTUs classifiable at different taxonomic ranks. Showing fractions of mapped reads, normalized per taxonomic rank. Eukaryotic OTUs were removed.

**Supplementary Figure 19:**
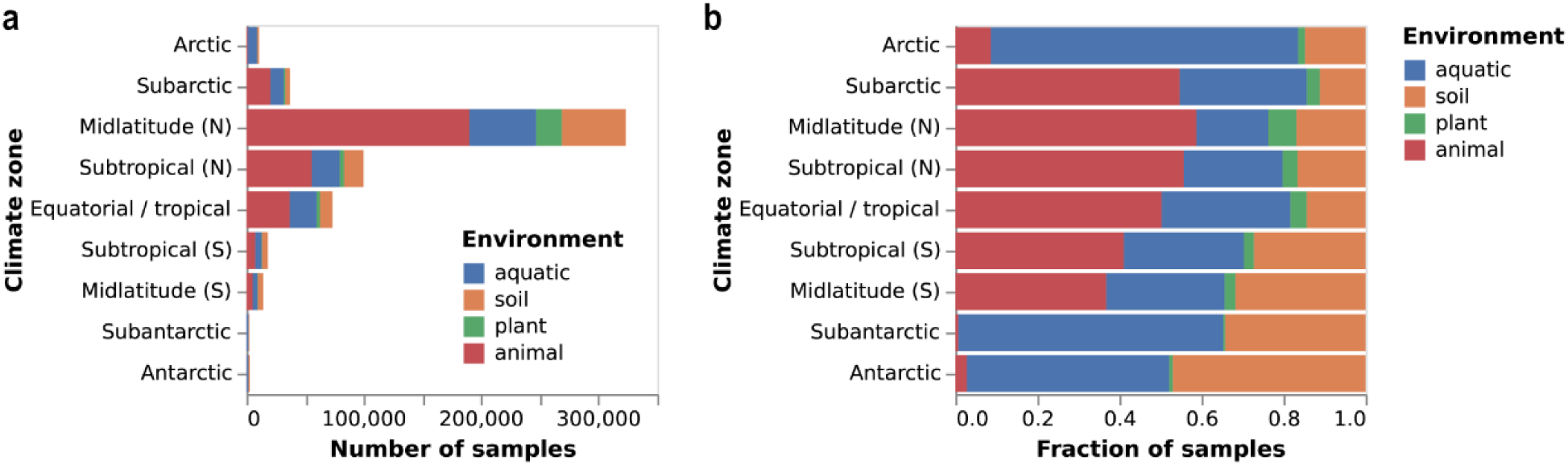
Distribution of MicrobeAtlas samples by environment and climate zone. Absolute number (a) and fraction (b) of samples per zone, stratified by MicrobeAtlas main environments.

**Supplementary Figure 20:**
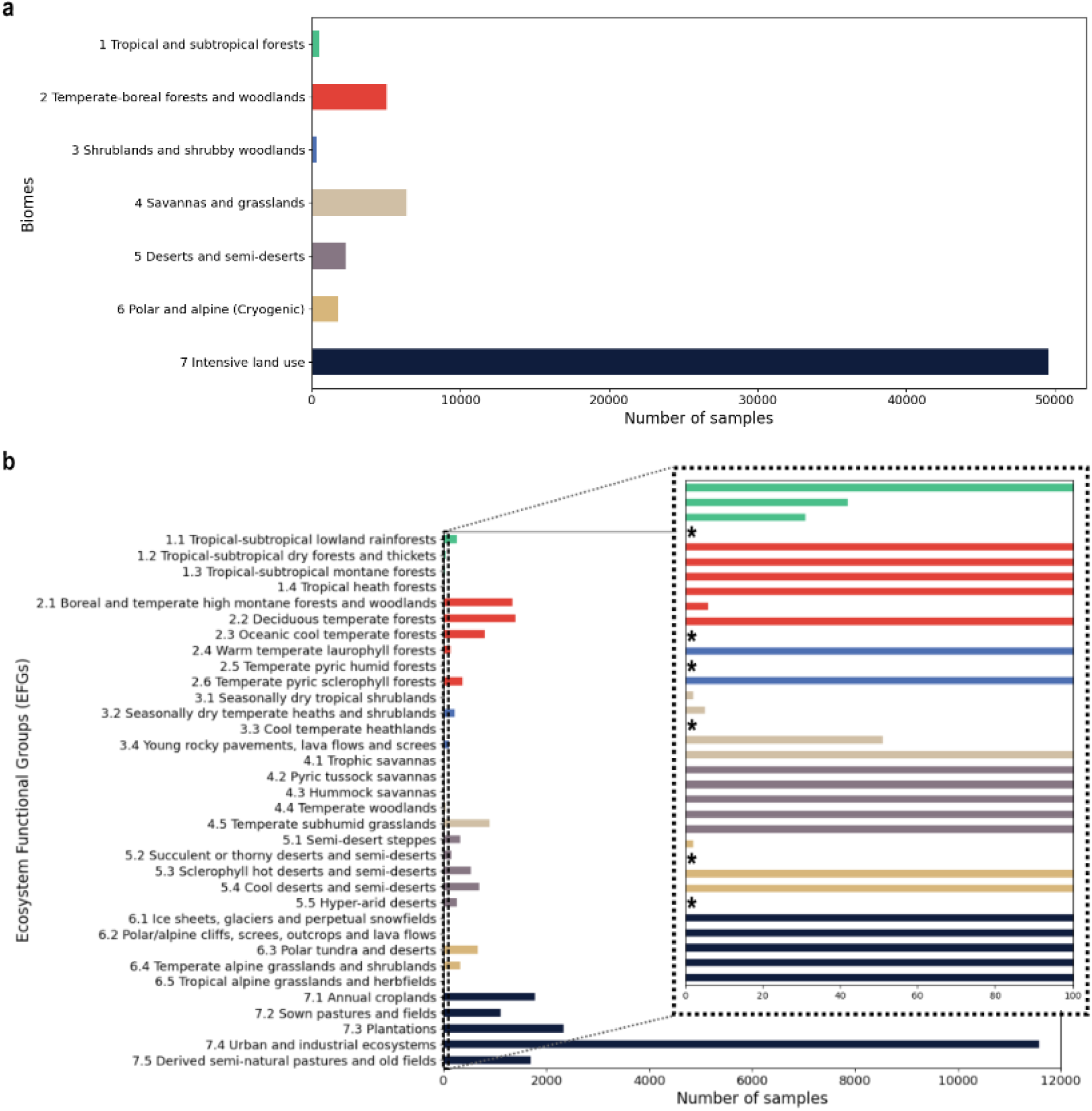
Number of samples across IUCN ecosystem definitions. Number of samples per IUCN Biome (a) and per IUCN Ecosystem Functional Groups (EFGs) (b) are shown. Asterisks in the zoom-in depict EFGs with no confidently geographically linked samples in MicrobeAtlas.

**Supplementary Figure 21:**
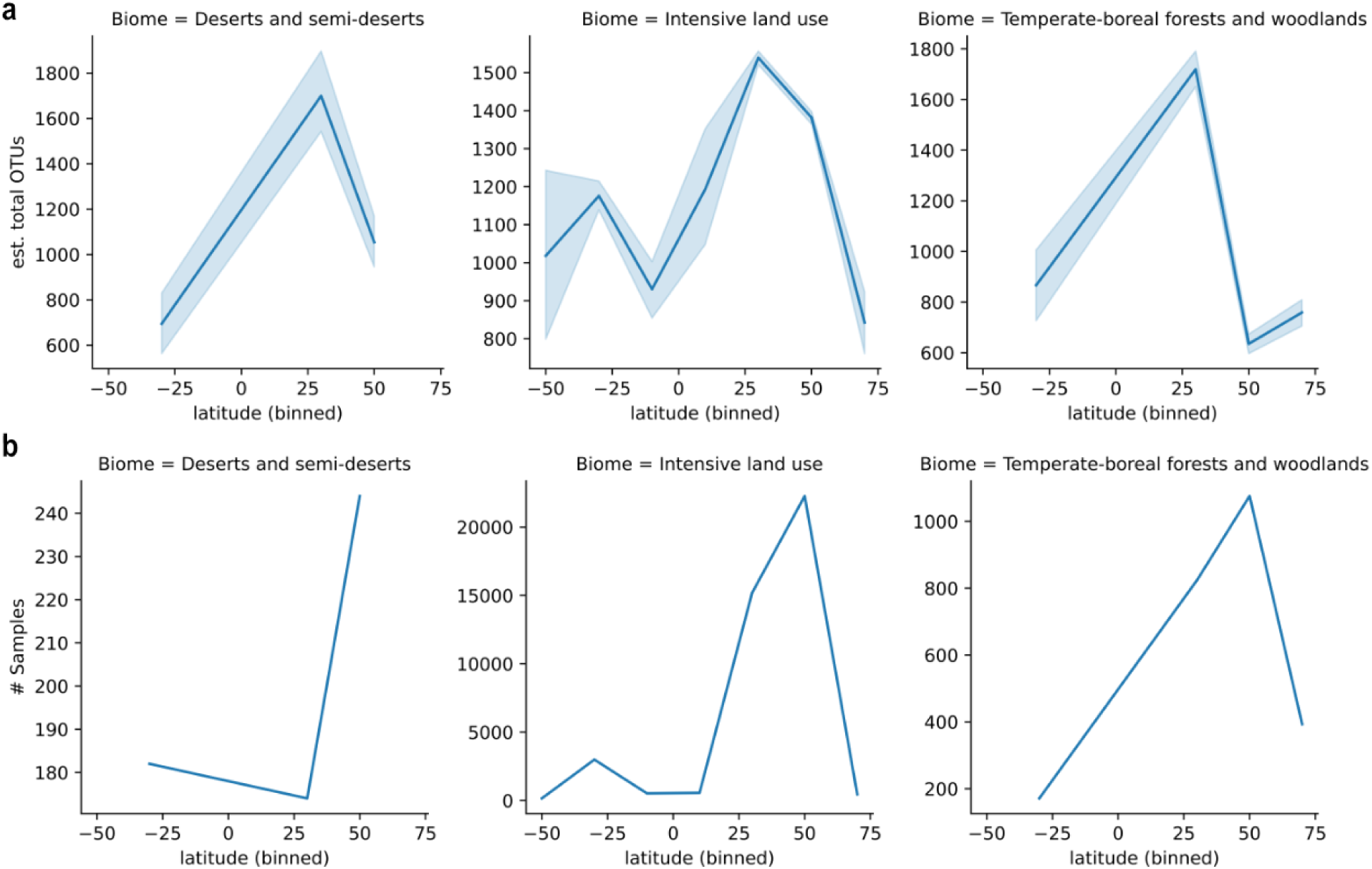
Latitudinal richness trends of terrestrial communities. a, Estimated richness of communities across latitudes for each robust IUCN biome (filtered as described in Methods). Lines and error bands represent means and 95% confidence intervals (1000 bootstraps) within 10 equally-sized latitudinal bins. **b**, Number of samples per latitudinal bin for each robust IUCN biome.

**Supplementary Figure 22:**
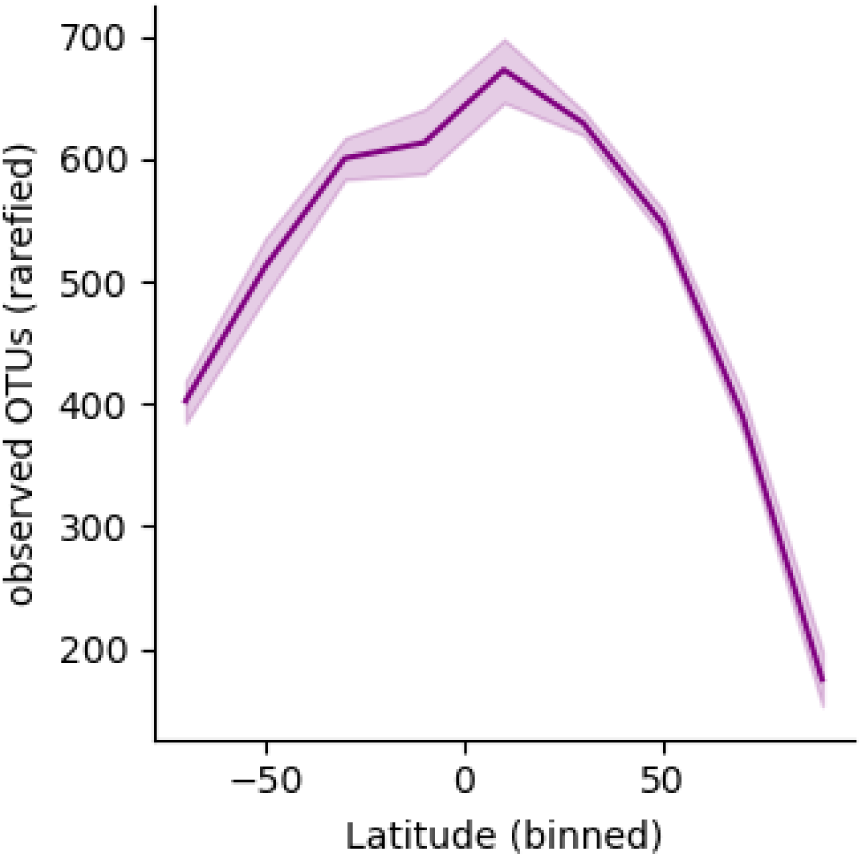
Observed rarefied OTU richness of marine communities across latitudes. Samples were uniformly rarefied to 1000 SSU reads. Lines and error bands represent means and 95% confidence intervals of log-transformed OTU counts (1000 bootstraps) within 10 equally-sized latitudinal bins.

**Supplementary Figure 23:**
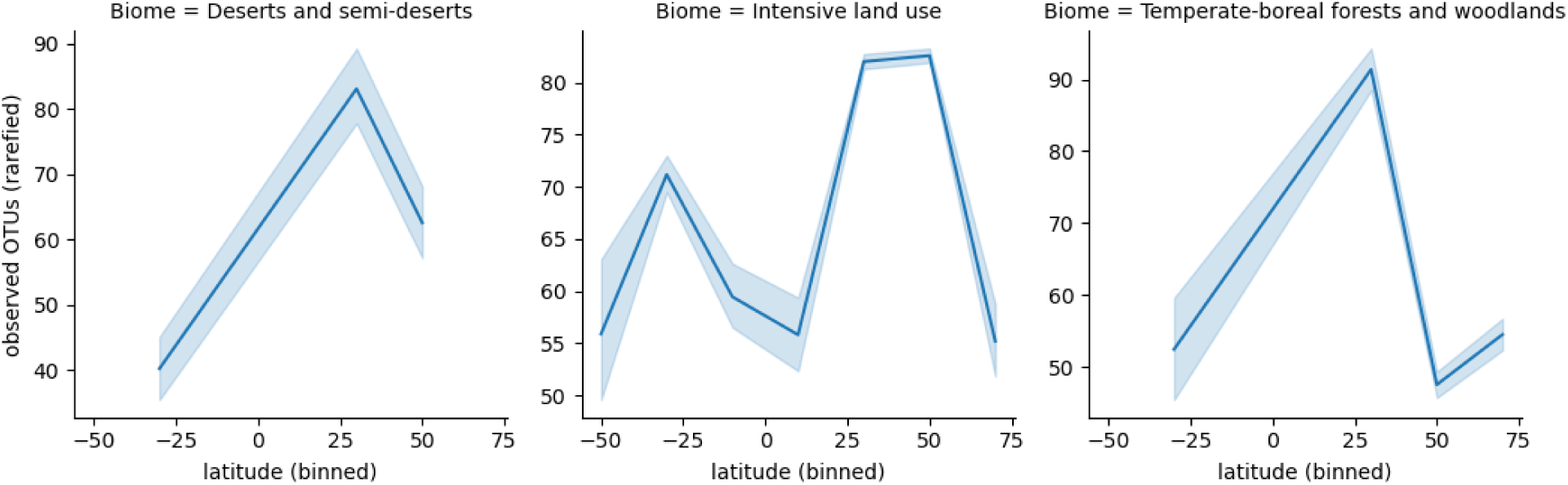
Observed latitudinal richness trends of terrestrial communities. a, Observed OTU richness (samples uniformly rarefied to 1000 SSU reads) of communities across latitudes for each robust IUCN biome (filtered as described in Methods). Lines and error bands represent means and 95% confidence intervals (1000 bootstraps) within 10 equally-sized latitudinal bins.

**Supplementary Figure 24:**
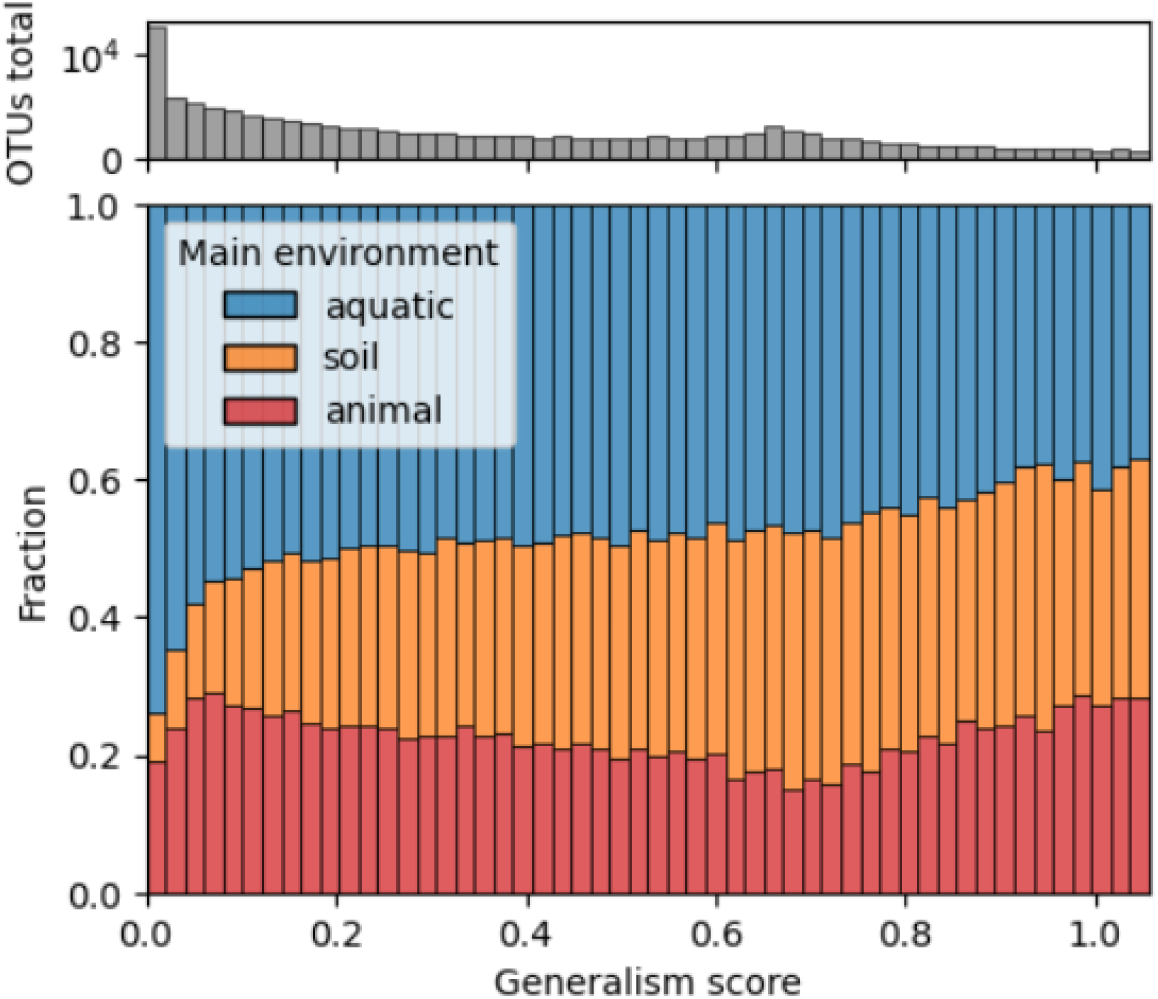
Number of OTUs and environmental preference shifts across the generalism spectrum. Top histogram: number of OTUs in each normalized generalism score bin. Bottom bar charts: fraction of OTUs in each generalism score bin, stratified by main environment (i.e. the environment in which each OTU exhibited its highest mean relative abundance). Generalism scores were computed excluding the “plant” environment.

**Supplementary Figure 25:**
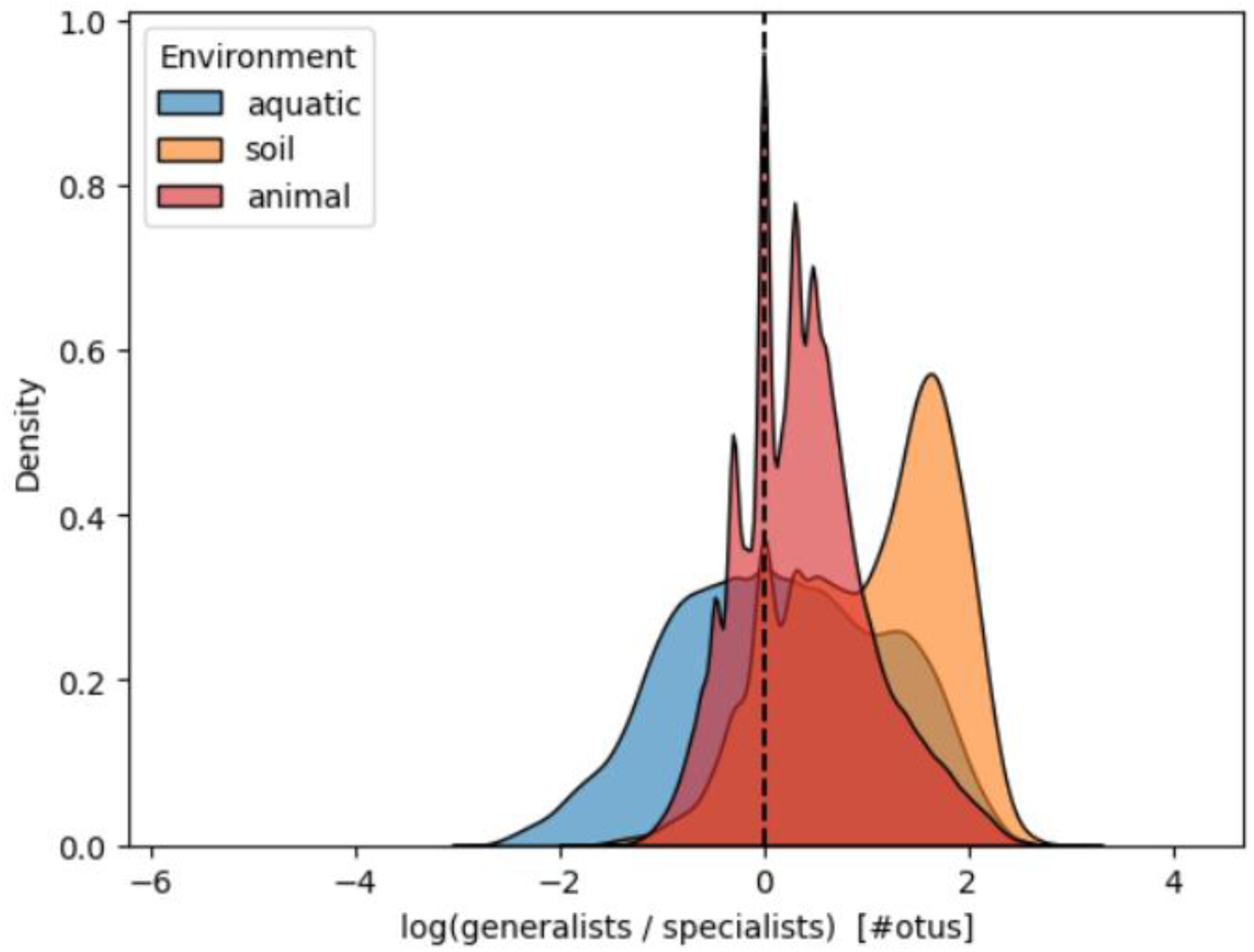
Generalist-to-specialist ratios across aquatic, soil, and animal samples. Log-ratios of the number of top-10% generalists to the number of top-10% specialists for each sample, grouped by environment. Generalist/specialist definitions were computed excluding the “plant” environment. Mean log-ratios: 1.03 (soil) vs. 0.32 (aquatic, animal), P < 10^-100^ (Mann-Whitney U test).

**Supplementary Figure 26:**
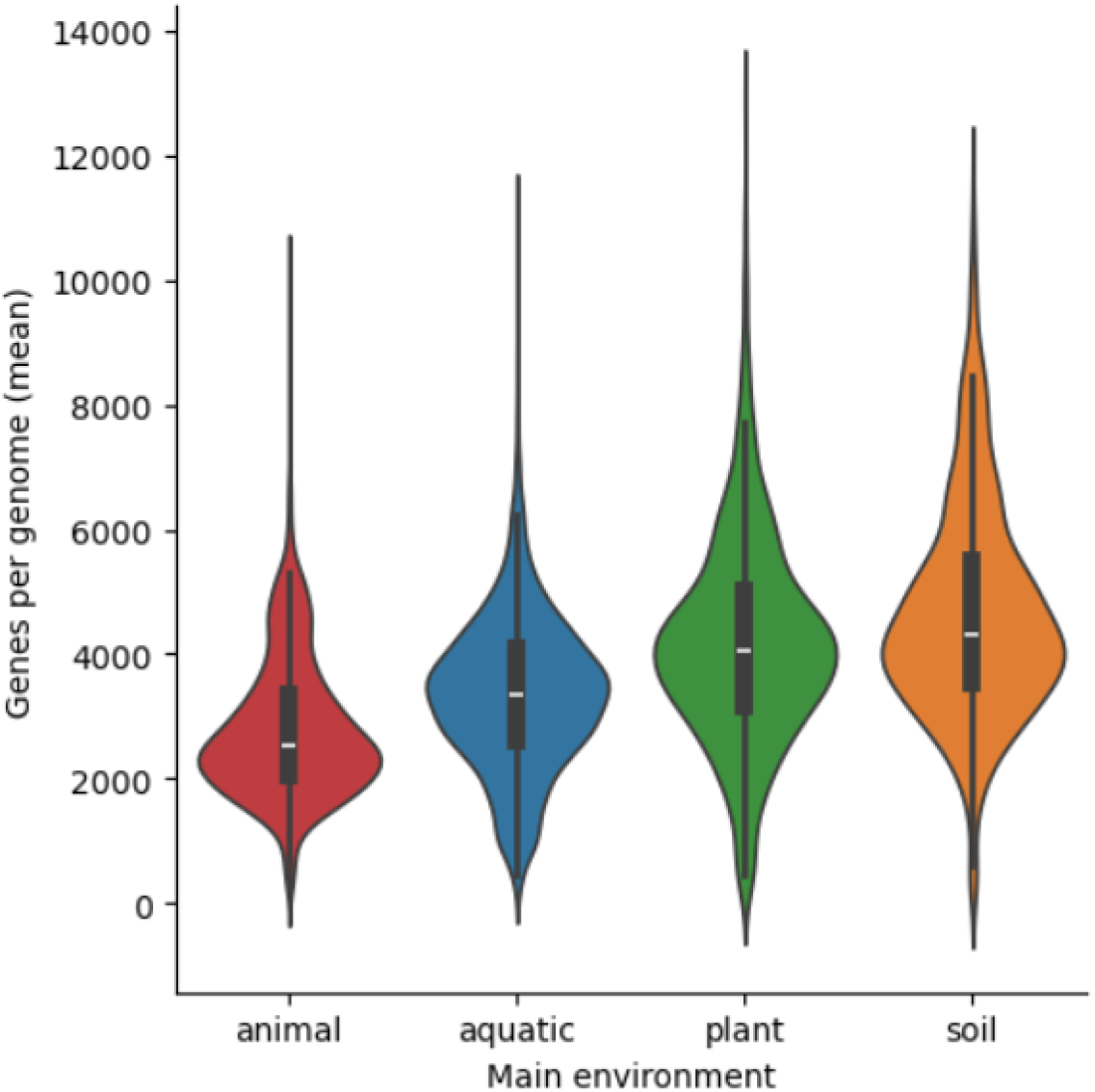
Genome sizes of 97%-level OTUs within their main environments. For each OTU with at least one mappable genome, the average gene count across its genomes is shown. OTUs sharing the same main environment (i.e. the environment in which each OTU exhibited its highest mean relative abundance) were grouped together. Boxes show interquartile ranges, whiskers represent 95% confidence intervals.

**Supplementary Figure 27:**
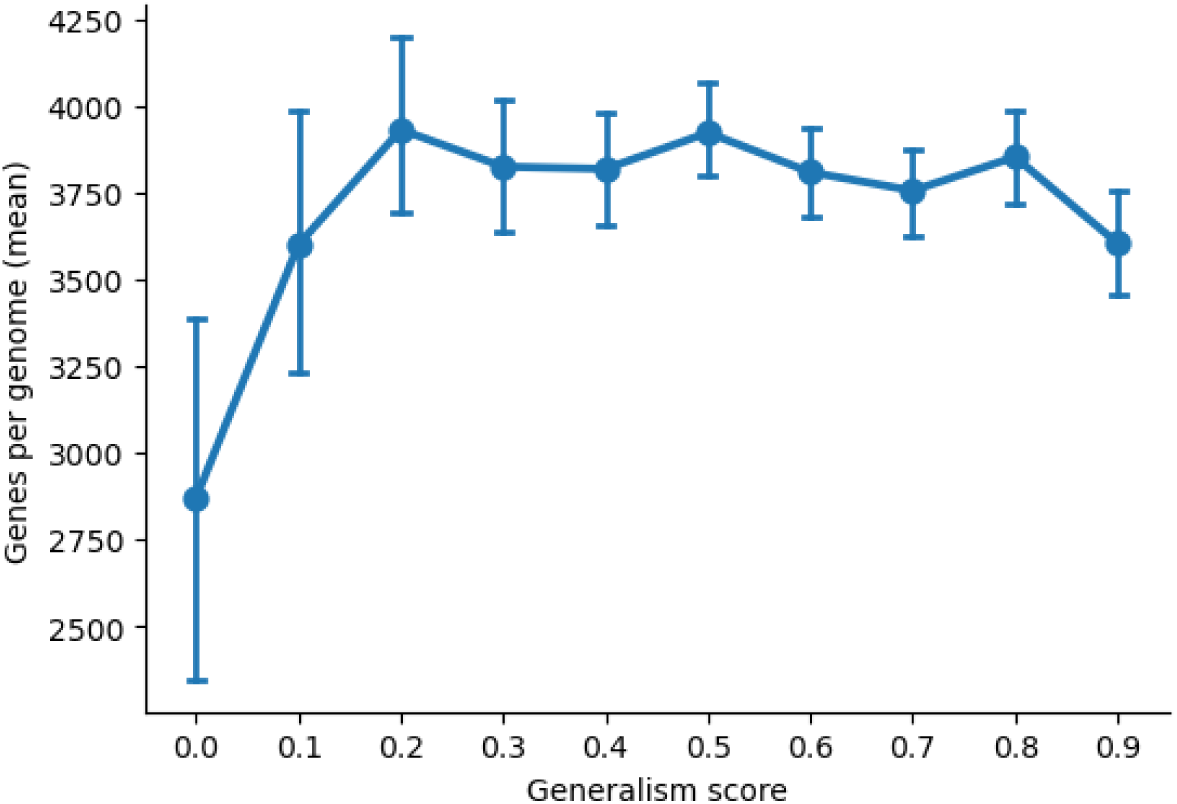
Relationships between genome size and generalism score, accounting for environmental distributions. Shows the average number of genes across proGenomes3-mappable OTUs within each generalism score bin. All generalism score bins were sub-sampled to represent all main environments (i.e. the environment in which each OTU exhibited its highest mean relative abundance) equally. Lines and error bands represent means and 95% confidence intervals (1000 bootstraps).

## Supplementary tables

**Supplementary Table 1:** The two-tiered environmental ontology used in MicrobeAtlas. Main and sub-environments assigned based on extracted, cleaned metadata keywords.

| Main environment | Sub environment | Main env. | Sub env. | Main env. | Sub env. |
| --- | --- | --- | --- | --- | --- |
| animal | baboon | animal (continued) | sparrow | aquatic (continued) | reservoir |
|  | bat |  | stork |  | river |
|  | bee |  | swallow |  | sea |
|  | bird |  | tadpole |  | sediment |
|  | bumblebee |  | termite |  | waste water |
|  | cat |  | tick | soil | agricultural |
|  | cattle |  | whale |  | desert |
|  | chimpanzee |  | zebrafish |  | farm |
|  | dog |  | roe |  | field |
|  | dolphin |  | seagull |  | forest |
|  | eagle |  | shark |  | paddy |
|  | fish |  | sheep |  | peatland |
|  | fly |  | pig |  | shrub |
|  | fruitfly |  | pigeon |  | tundra |
|  | goat |  | rat | plant | flower |
|  | gorilla | aquatic | brine |  | leaf |
|  | hamster |  | estuary |  | rhizosphere |
|  | horse |  | groundwater |  | seed |
|  | human |  | ice |  | sprout |
|  | macaque |  | lake |  | stem |
|  | mosquito |  | marine |  | wood |
|  | mouse |  | ocean |  |  |

**Supplementary Table 2:** Number of samples per IUCN Biome. Only samples that could be confidently spatially mapped to a single IUCN Biome are reported.

| IUCN Biome | Number of samples |
| --- | --- |
| 1 Tropical and subtropical forests | 516 |
| 2 Temperate-boreal forests and woodlands | 5042 |
| 3 Shrublands and shrubby woodlands | 350 |
| 4 Savannas and grasslands | 6373 |
| 5 Deserts and semi-deserts | 2260 |
| 6 Polar and alpine (Cryogenic) | 1758 |
| 7 Intensive land use | 49610 |

**Supplementary Table 3:**
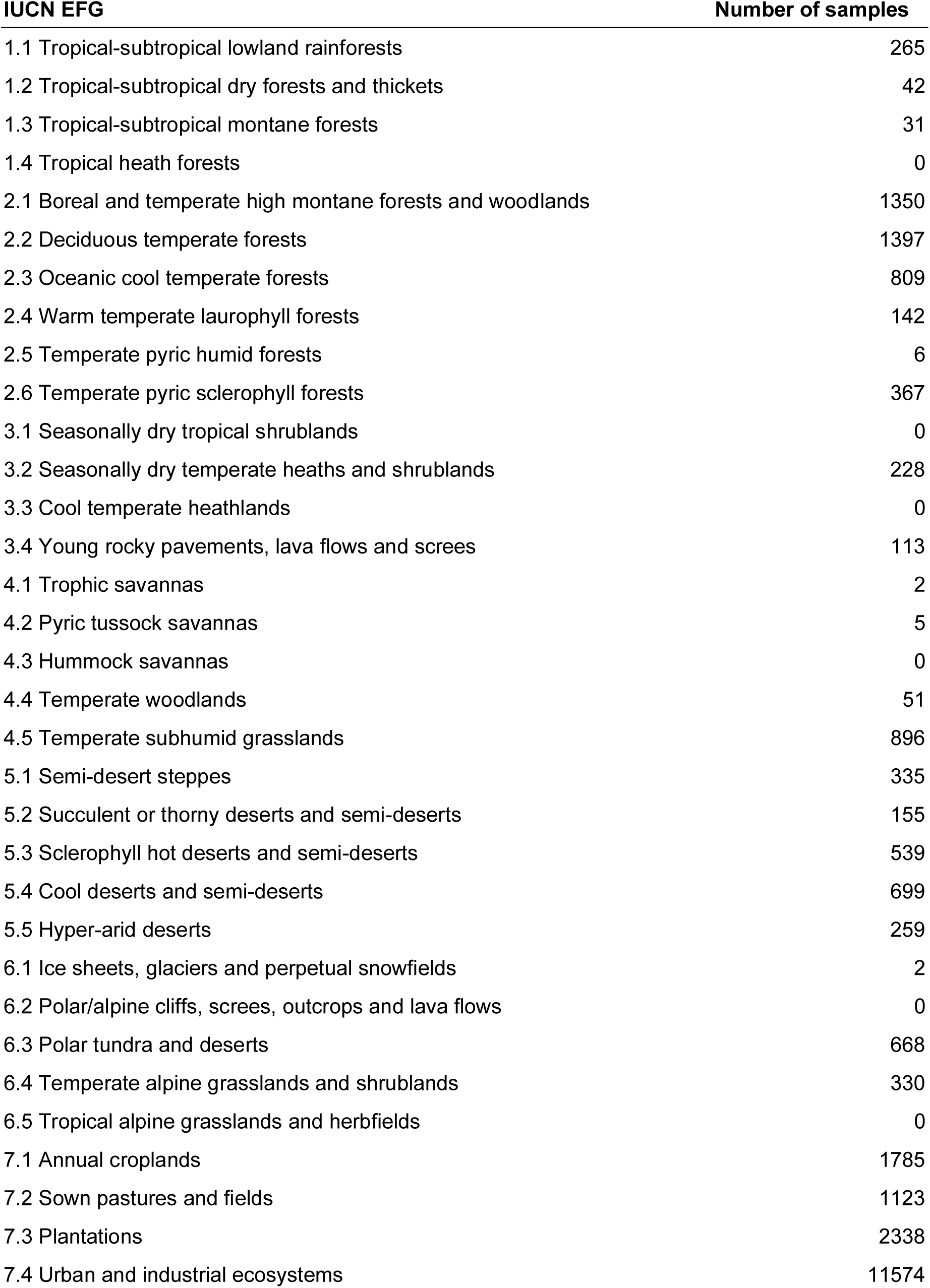
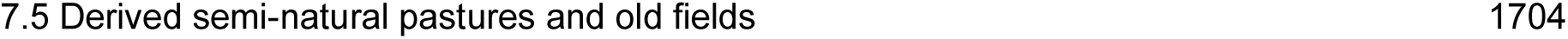
Number of samples per IUCN Ecosystem Functional Group (EFG). Only samples that could be confidently spatially mapped to a single IUCN EFG are reported.

| **IUCN EFG** | **Number of samples** |
| --- | --- |
| 1.1 Tropical-subtropical lowland rainforests | 265 |
| 1.2 Tropical-subtropical dry forests and thickets | 42 |
| 1.3 Tropical-subtropical montane forests | 31 |
| 1.4 Tropical heath forests | 0 |
| 2.1 Boreal and temperate high montane forests and woodlands | 1350 |
| 2.2 Deciduous temperate forests | 1397 |
| 2.3 Oceanic cool temperate forests | 809 |
| 2.4 Warm temperate laurophyll forests | 142 |
| 2.5 Temperate pyric humid forests | 6 |
| 2.6 Temperate pyric sclerophyll forests | 367 |
| 3.1 Seasonally dry tropical shrublands | 0 |
| 3.2 Seasonally dry temperate heaths and shrublands | 228 |
| 3.3 Cool temperate heathlands | 0 |
| 3.4 Young rocky pavements, lava flows and screes | 113 |
| 4.1 Trophic savannas | 2 |
| 4.2 Pyric tussock savannas | 5 |
| 4.3 Hummock savannas | 0 |
| 4.4 Temperate woodlands | 51 |
| 4.5 Temperate subhumid grasslands | 896 |
| 5.1 Semi-desert steppes | 335 |
| 5.2 Succulent or thorny deserts and semi-deserts | 155 |
| 5.3 Sclerophyll hot deserts and semi-deserts | 539 |
| 5.4 Cool deserts and semi-deserts | 699 |
| 5.5 Hyper-arid deserts | 259 |
| 6.1 Ice sheets, glaciers and perpetual snowfields | 2 |
| 6.2 Polar/alpine cliffs, screes, outcrops and lava flows | 0 |
| 6.3 Polar tundra and deserts | 668 |
| 6.4 Temperate alpine grasslands and shrublands | 330 |
| 6.5 Tropical alpine grasslands and herbfields | 0 |
| 7.1 Annual croplands | 1785 |
| 7.2 Sown pastures and fields | 1123 |
| 7.3 Plantations | 2338 |
| 7.4 Urban and industrial ecosystems | 11574 |

**Supplementary Table 4:** Semantic keyword clusters enriched in mice compared to human communities. Semantic clusters are based on large language model semantic clustering (Methods). Mean odds ratio represents the average fold change in prevalence of a given keyword within a cluster, comparing mouse to human. Mean counts represent the average number of times terms from each semantic cluster appeared in mouse samples. Only the top 15 enriched clusters are shown.

| Semantic cluster | Mean odds ratio | Mean count (mouse) |
| --- | --- | --- |
| host,intergenerational,outbred,commensalism,<br>symbiosis,homozygous,multigenerational | 1.50 | 2224.00 |
| exposure,disturbance,dose-dependent,disruption,<br>resistance,transmission | 1.50 | 1475.33 |
| innate,epigenome,autologous | 1.50 | 706.67 |
| toxicity,irradiation,radiation,heat-killed,redox,<br>radioactivity | 1.50 | 585.67 |
| anatomy,skin,tissue,organ,musculus,brain,bone | 1.50 | 503.57 |
| insulin,hormone | 1.50 | 356.50 |
| gavaged,goblet,tract,swab,distal,lumen | 1.42 | 700.83 |
| animal,mouse,rat,rodent,mammal,pup,vole,bird,dog | 1.39 | 6149.78 |
| lung,epithelium,pharynx,vagina,mucosa,liver,<br>urogenital tract,mucin,blood | 1.39 | 649.22 |
| deficiency,immunity,stress,impaired,anti-inflammatory,<br>immunization,susceptibility,dysfunction | 1.38 | 994.00 |
| induction,dosage,removal,synthesis,reduction,<br>reconstitution,intake,supplementation | 1.38 | 905.13 |
| acarbose,linolenic,linoleic,fermentation | 1.38 | 281.50 |
| antibiotic,antibiotic-perturbed,anti-bacterial,rifaximin,<br>streptomycin,antimicrobial,antibiotic-induced,<br>antibiotic-treated,post-antibiotic,vancomycin | 1.35 | 1033.10 |
| human-gut,animal-associated,gnotobiotic,polygyrus,<br>heligmosomoides,xenomicrobiota,mitochondria,<br>flagellin,mouse-associated,murine-feces,musculus-<br>gut-bacteria,germfree | 1.33 | 928.17 |
| perturbed,degradation,inhibition,attenuation,etiology,<br>perturbation | 1.33 | 454.33 |

**Supplementary Table 5:**
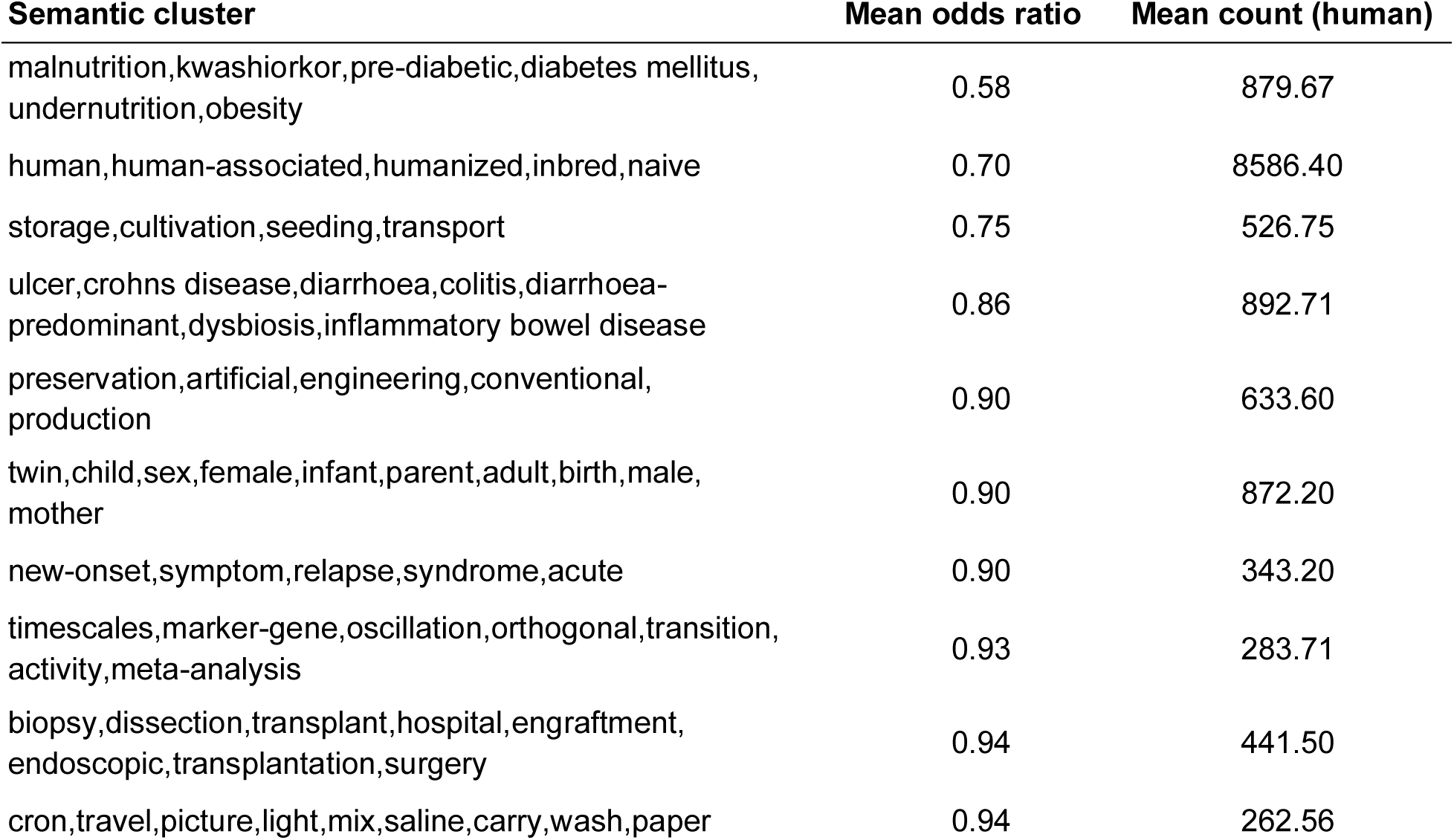
Semantic keyword clusters depleted in mice compared to human communities. Semantic clusters are based on large language model semantic clustering (Methods). Mean odds ratio represents the average fold change in prevalence of a given keyword within a cluster, comparing mouse to human. Mean counts represent the average number of times terms from each semantic cluster appeared in human samples. Only the top 15 depleted clusters are shown.

| Semantic cluster | Mean odds ratio | Mean count (human) |
| --- | --- | --- |
| malnutrition,kwashiorkor,pre-diabetic,diabetes mellitus, undernutrition,obesity | 0.58 | 879.67 |
| human,human-associated,humanized,inbred,naive | 0.70 | 8586.40 |
| storage,cultivation,seeding,transport | 0.75 | 526.75 |
| ulcer,crohns disease,diarrhoea,colitis,diarrhoea-predominant,dysbiosis,inflammatory bowel disease | 0.86 | 892.71 |
| preservation,artificial,engineering,conventional, production | 0.90 | 633.60 |
| twin,child,sex,female,infant,parent,adult,birth,male, mother | 0.90 | 872.20 |
| new-onset,symptom,relapse,syndrome,acute | 0.90 | 343.20 |
| timescales,marker-gene,oscillation,orthogonal,transition, activity,meta-analysis | 0.93 | 283.71 |
| biopsy,dissection,transplant,hospital,engraftment, endoscopic,transplantation,surgery | 0.94 | 441.50 |
| cron,travel,picture,light,mix,saline,carry,wash,paper | 0.94 | 262.56 |

**Supplementary Table 6: The top 150 most abundant OTUs within MicrobeAtlas.** Mean relative abundances across all samples were used to rank OTUs. Top OTUs were further annotated with taxonomic assignments and OTU type (cg-OTU or ncg-OTU).

**Supplementary Table 7: Metadata keywords indicative of each main environment.** Terms used to assign each sample to one or more main environments (if possible).

